# Insights into Dentatorubral-Pallidoluysian Atrophy from a new *Drosophila* model of disease

**DOI:** 10.1101/2024.12.05.627083

**Authors:** Matthew V. Prifti, Oluwademilade Nuga, Ryan O. Dulay, Nikhil C. Patel, Truman Kula, Kozeta Libohova, Autumn Jackson-Butler, Wei-Ling Tsou, Kristin Richardson, Sokol V. Todi

**Affiliations:** Department of Pharmacology, Wayne State University School of Medicine; Department of Neurology, Wayne State University School of Medicine

**Keywords:** ataxia, atrophin, dementia, myoclonus, neurodegeneration, polyglutamine, protein folding, quality control

## Abstract

Dentatorubral-pallidoluysian atrophy (DRPLA) is a neurodegenerative disorder that presents with ataxia, dementia and epilepsy. As a member of the polyglutamine family of diseases, DRPLA is caused by abnormal CAG triplet expansion beyond 48 repeats in the protein-coding region of *ATROPHIN 1* (*ATN1*), a transcriptional co-repressor. To better understand DRPLA, we generated new *Drosophila* lines that express full-length, human ATN1 with a normal (Q7) or pathogenic (Q88) repeat. Expression of ATN1 is toxic, with the polyglutamine-expanded version being consistently more problematic than wild-type ATN1. Fly motility, longevity and internal structures are negatively impacted by pathogenic ATN1. RNA-seq identified altered protein quality control and immune pathways in the presence of pathogenic ATN1. Based on these data, we conducted genetic experiments that confirmed the role of protein quality control components that ameliorate or exacerbate ATN1 toxicity. Hsc70-3, a chaperone, arose as a likely suppressor of toxicity. VCP (a proteasome-related AAA ATPase), Rpn11 (a proteasome-related deubiquitinase) and select DnaJ proteins (co-chaperones) were inconsistently protective, depending on the tissues where they were expressed. Lastly, informed by RNA-seq data that exercise-related genes may also be involved in this model of DRPLA, we conducted short-term exercise, which improved overall fly motility. This new model of DRPLA will prove important to understanding this understudied disease and will help to identify therapeutic targets for it.

## INTRODUCTION

The polyglutamine (polyQ) family of disorders consists of nine inherited, age-related neurodegenerative diseases including Huntington’s Disease (HD), Spinal and Bulbar Muscular Atrophy (SBMA, also known as Kennedy’s Disease), Spinocerebellar Ataxias (SCA) type 1, 2, 3, 6, 7 and 17, and Dentatorubral-Pallidoluysian Atrophy (DRPLA)^1–8^. DRPLA, the focus of this report, impacts a small number of individuals worldwide, with affected population clusters found mainly in Japan and with lower incidence being described in Europe^9^. The pathology presents clinically with motor and cognitive symptoms including ataxia, cognitive decline and epilepsy^10,11^. Degeneration in this disease mainly occurs in the dentatorubral and pallidoluysian systems, which are collectively involved in the coordination and regulation of movement. Besides SBMA, which is X-linked, all polyQ disorders are inherited in an autosomal dominant manner^1,3,7^; they display clinically distinct pathologies and are caused by the abnormal expansion of the polyQ tract within the affected protein. In DRPLA, this expansion (>49 repeats) occurs within the protein Atrophin-1 (ATN1), which is encoded by the *ATROPHIN-1* (*ATN1*) gene on chromosome 12^4^. Like the other diseases of the polyQ family, patients with DRPLA show an inverse relationship between repeat length and age of onset^1,3,12^.

Atrophins are transcriptional regulators conserved from invertebrates to mammals. In humans, there are two *ATROPHIN* genes: *ATROPHIN-1* and *ATROPHIN-2* (a.k.a. RERE)^13^. Both proteins have been implicated in development across organisms, specifically for correct segmentation and embryonic patterning in *Drosophila* and embryogenesis, neurodevelopment and cardiac development in mice^14–18^. Mutations in the non-polyQ region of exon 7 in *ATN1* cause ATN1-related neurodevelopmental disorder (ATN1-NDD), a disease characterized by cognitive deficits, reduced muscle tone and malformations of the hands and feet^19^. ATN1 is thought to mainly function as a transcriptional co-repressor^16^. When polyQ-expanded in DRPLA, ATN1 seems to acquire properties that may lead to toxicity^1,3,20^. These include reduced phosphorylation that may delay neuronal cell survival pathway activation^21^, impaired autophagy preventing elimination of polyQ protein^22^ and decreased protein stability leading to abnormal cleavage or fragmentation and the formation of nuclear inclusions^23^. Still, the biology of disease of DRPLA remains unclear and, as with other disorders of the polyQ family, no therapeutic options are available.

Towards better understanding the DRPLA biology of disease and finding therapeutic options for it, we generated DRPLA models in *Drosophila melanogaster.* The fruit fly has a long history of impactful use towards understanding function and diseases of the nervous system^24–26^. Here, we describe the generation and initial characterization of isogenic fly lines expressing full-length, human ATN1 with a wild-type (Q7) or polyQ-expanded (Q88) repeat that is within the human patient range^1^. We found variable toxicity with both wild-type (WT) and disease-causing (DRPLA) ATN1 expression when measuring lifespan, depending on the tissue where these genes were expressed, as well as impairments in motility and endurance. On a cellular level, we observed accumulation of ATN1 protein over time. RNA-seq analyses in combination with genetic interrogation identified several protein quality control and immune pathway constituents whose targeting enhanced or suppressed toxicity. Alongside various chaperones and proteasome-related components that we identified as potential therapeutic targets for DRPLA, endurance exercise also emerged as a suppressor. Collectively, the data in this report highlight the new *Drosophila* models of DRPLA as important tools to understand this disease and to unravel the contribution of key genes and proteins in its pathology and future treatment.

## RESULTS

### Expression of ATN1 is toxic in *Drosophila*

We previously reported the generation of various *Drosophila* models of polyQ disorders: SBMA, SCAs 3, 6, 7 and 17^5,27–35^, as well as additional lines that are undergoing thorough characterization. For each, we used phiC31-dependent integration^36,37^ to incorporate the full-length human genes into a “safe-harbor” on chromosome 3, making them comparable to one another and to their WT controls by eliminating effects of random insertion, ensuring the incorporation of a single transgene in the same orientation and utilizing the same vector backbone for all transgenes (pWalium10.moe). This toolbox of polyQ fly models now includes DRPLA.

To generate ATN1 transgenic flies, we synthesized cDNA sequences encoding full-length, human ANT1 with either a non-disease (CAG/CAA-Q7) or DRPLA-causing (CAG/CAA-Q88) polyQ (figure 1A) reflecting the upper end of repeats seen in DRPLA patients^1^. We added a terminal HA epitope tag to both versions. The transgenes were inserted into the attP2 site on chromosome 3. Like the other polyQ disease models in the fly, the DRPLA lines are designed to be used with the Gal4-UAS binary system of expression^38^. To control for potential non-traditional (non-AUG) protein translation or unpredicted mRNA toxicity because of expanded CAG repeats, we designed the polyQ tracts to be encoded by alternating CAG/CAA repeats. We have used this design of the polyQ tract in our other polyQ *Drosophila* models^5,27,28^. To confirm the transgenes’ presence, we conducted PCR analyses of the attP2 insertion sites of WT and polyQ-expanded ATN1, which confirmed integration into attP2 (figure 1B). Genomic DNA sequence analyses confirmed the identity and integrity of the intended transgenes (data not included).

**Figure 1:**
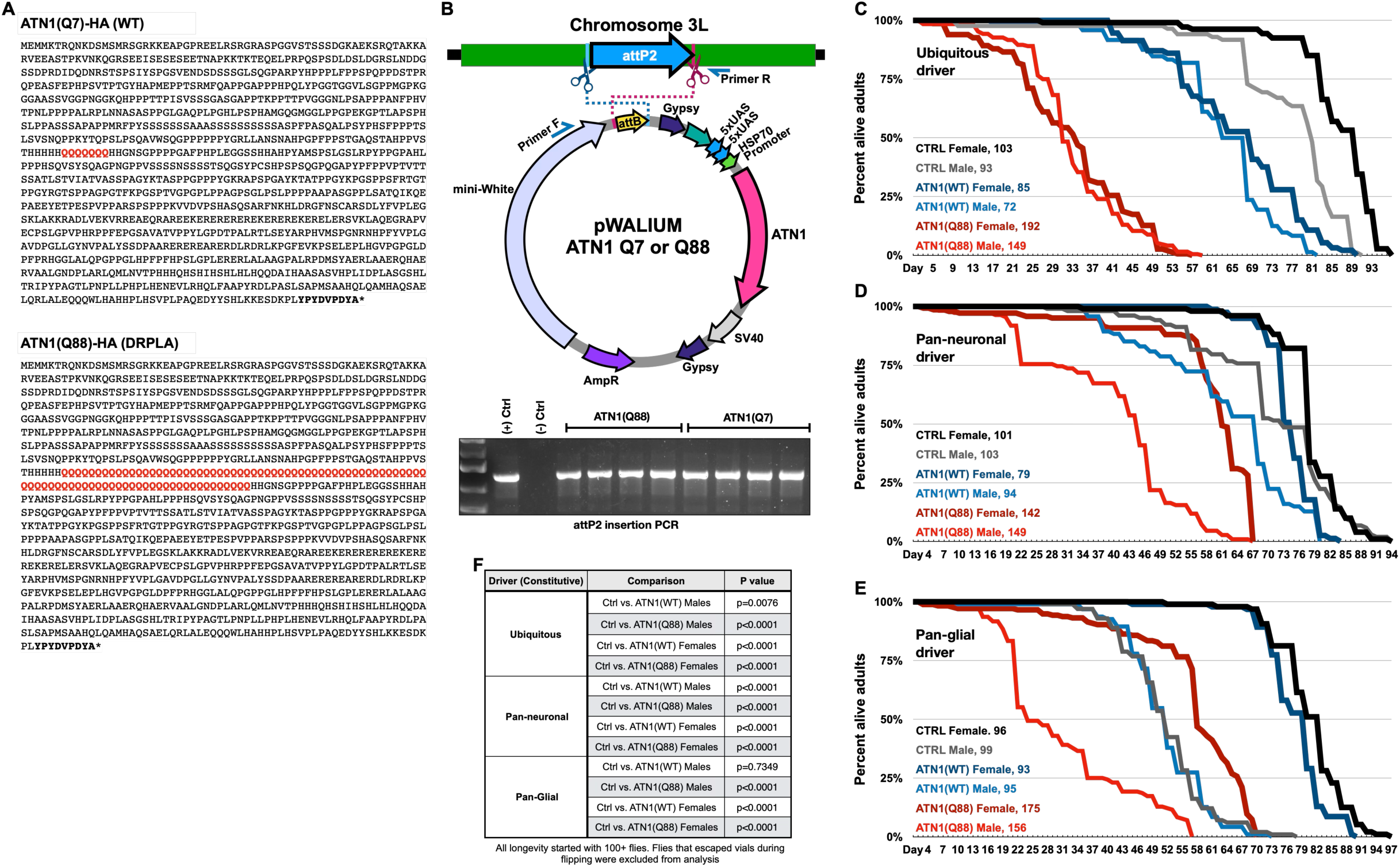
Effect of ATN1 protein expression on fly longevity. A) Amino acid used for ATN1(WT) (Q7, top) and ATN1(Q88) (bottom). The polyQ tract is in red font. The C-Terminal HA tag is emphasized in bold typeface. B) Schematic representation of the cloning plasmid used for ATN1 cDNA integration into the pWallium10.moe vector (top). Confirmation of the insertion sites in the created lines (bottom). C-E) Longevity curves for ATN1(Q88) (red), ATN1(WT) (blue), and CTRL (black, gray) with (C) ubiquitous, (D) pan-neuronal and (E) pan-glial expression. F) Statistical analyses for longevity curves from panels C-E: Log-Rank tests ‘N’s are indicated in the figure panels.

We utilized the Gal4-UAS system to express WT (Q7) or disease-causing (DRPLA; Q88) ATN1 in fly tissues. We used the following three expression patterns: ubiquitous (driven by sqh-Gal4), pan-neuronal (elav-Gal4) and pan-glial (Repo-Gal4). Each of these drivers leads to transgene expression during development and continuing through adulthood. We selected these patterns since the *ATN1* gene is widely expressed and since the nervous system is especially perturbed in all polyQ disorders, including in DRPLA. We found that expression of either form of ATN1 in all tissues was toxic compared to control flies that contained the sqh-Gal4 driver in the same genetic background as the ATN1 lines (w^1118^; figure 1C). Expression of the pathogenic form was markedly more toxic than WT. The fact that over-expression of a WT polyQ disease protein is problematic in the fly is not surprising, as it has been described before for other polyQ disease models and likely results from interference of the expressed transgene with endogenous functions^27,28^. Sex-dependent differences were apparent in the Ctrl and WT groups, but not in the DRPLA groups in this expression pattern. Pan-neuronal expression of ATN1 was also toxic, especially for the disease-causing form and particularly in male flies (figure 1D). Expression of the WT variant was non-toxic in females and mildly toxic in males, underscoring sex-dependent effects (figure 1D). With pan-glial expression, we observed that the disease-causing version of ATN1 was toxic to female and especially male flies, whereas the WT version was well tolerated by both sexes (figure 1E; statistical comparisons for the longevity examinations compared to control flies are summarized in figure 1F). Collectively, longevity assays indicate toxicity from the Q88 version of ATN1, with mild to no toxicity from Q7 ATN1.

Because DRPLA affects motor functions, we next examined adult fly motility when the WT and DRPLA versions of ATN1 were expressed in all neurons or glia (figure 2A). With pan-neuronal expression, we observed a mild but statistically significant reduction in fly mobility over time when the DRPLA form was present in male and female flies. Expression of WT ATN1 did not lead to reduced mobility – in fact, it led to increased performance in both female and male flies at select time points (e.g., day 1, week 4). Expression in all glia also led to a mild decrease in mobility in both sexes over time. With this expression pattern, WT ATN1 again led to increased mobility at some time points or to no difference from control flies in both sexes. Collectively, these results indicate mild reduction in motility from flies expressing disease-causing ATN1.

**Figure 2:**
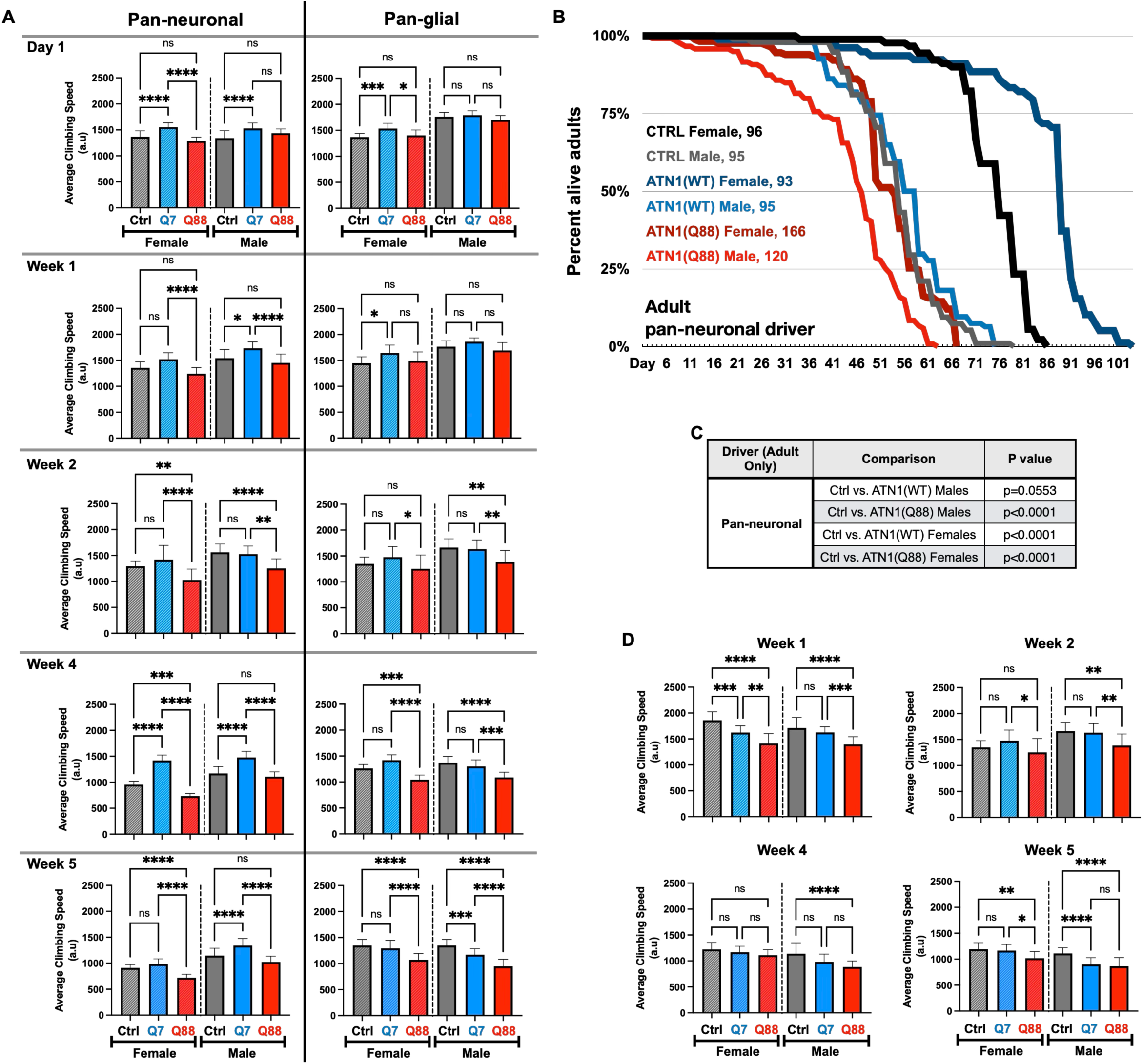
Effect of ATN1 protein expression on fly motility. A) Average climbing index for Ctrl (gray), ATN1(WT) (blue) and ATN1(Q88) (red) expression in neurons and in glia. One-way ANOVA + Tukey post-hoc. N=100 per cohort. B) Longevity curves from adult, pan-neuronal expression of ATN1(WT) (blue), ATN1(Q88) (red), and non-expressing controls (black, grey). C) Statistical analyses used for (B): Log-Rank tests, N≥100 per cohort. D) Average climbing index following adult pan-neuronal expression of Ctrl (gray), ATN1(WT) (blue) and ATN1(Q88) (red). One-way ANOVA + Tukey post-hoc. N=100 per cohort. For all motility graphs in panels (A) and (D): *: p<0.05, **: p<0.005, ***: p<0.001, ****: p<0.0001.

Because DRPLA is largely adult-onset, we next examined the toxicity of WT and pathogenic ATN1 in flies that express these transgenes only in neurons, starting as day 1-adults. For this approach, we used RU486-dependent expression of ATN1 through the GeneSwitch (GS) Gal4-UAS system^39^. In this system, RU486 is required to activate expression. Our driver was the pan-neuronal elavGS-Gal4. We reared flies in media without RU486 until the day that they eclosed as adults, when they were switched to media containing the inducer. Our findings are summarized in figures 2B-D. We found that pathogenic ATN1 was toxic in both sexes, whereas WT ATN1 had no effect in male flies and extended longevity in female flies (figure 2B, 2C). In terms of motility, we again found mild reduction in performance from flies expressing pathogenic ATN1, with little or no effect from the WT version over time (figure 2D). Based on the collective results of figure 2, we conclude that pathogenic ATN1 is consistently toxic in *Drosophila*, whereas the WT version is generally not problematic.

### Human ATN1 protein in fly tissues

We conducted Western blotting (WB) of ATN1 expressed in fly tissues to examine its migration pattern. Figure 3 summarizes our findings from constitutive ubiquitous (figure 3A) and adult pan-neuronal (figure 3B) expression. We examined both males and females. The WT and DRPLA versions of ATN1 migrate as expected, with the pathogenic species migrating more slowly on SDS-PAGE gels. Besides the main bands highlighted by arrows in figure 3, we also observed various, lower molecular weight species that likely correspond to processed ATN1 by proteases. ATN1 can be proteolyzed into smaller fragments in cells^1,23^. With ubiquitous expression, we observed a trend of reduced intensity of the primary band of ATN1 as time progressed in both the WT and Q88 variants (figure 3A; the likely proteolytic bands show a similar tendency). This is likely a result of aggregation of both the WT and DRPLA versions of ATN1 over time; such species are then pelleted during the centrifugation step of homogenate preparation for WB and are thus excluded from SDS-PAGE gels (Methods). To address this possibility, we conducted modified centrifugation protocols to separate ATN1 protein into soluble and pellet fractions. As shown in supplemental figure 1, we observed increased levels of ATN1(Q88) in the pellet over time; ATN1(Q7) is also consistently present in the pellet at all times investigated. Additionally, we also noted punctate staining in dissected fly brains stained for ATN1, which appears to become visibly more pronounced at aged time-points (supplemental figure 2). Based on these results, both WT and disease-causing ATN1 aggregate in the fly.

**Figure 3:**
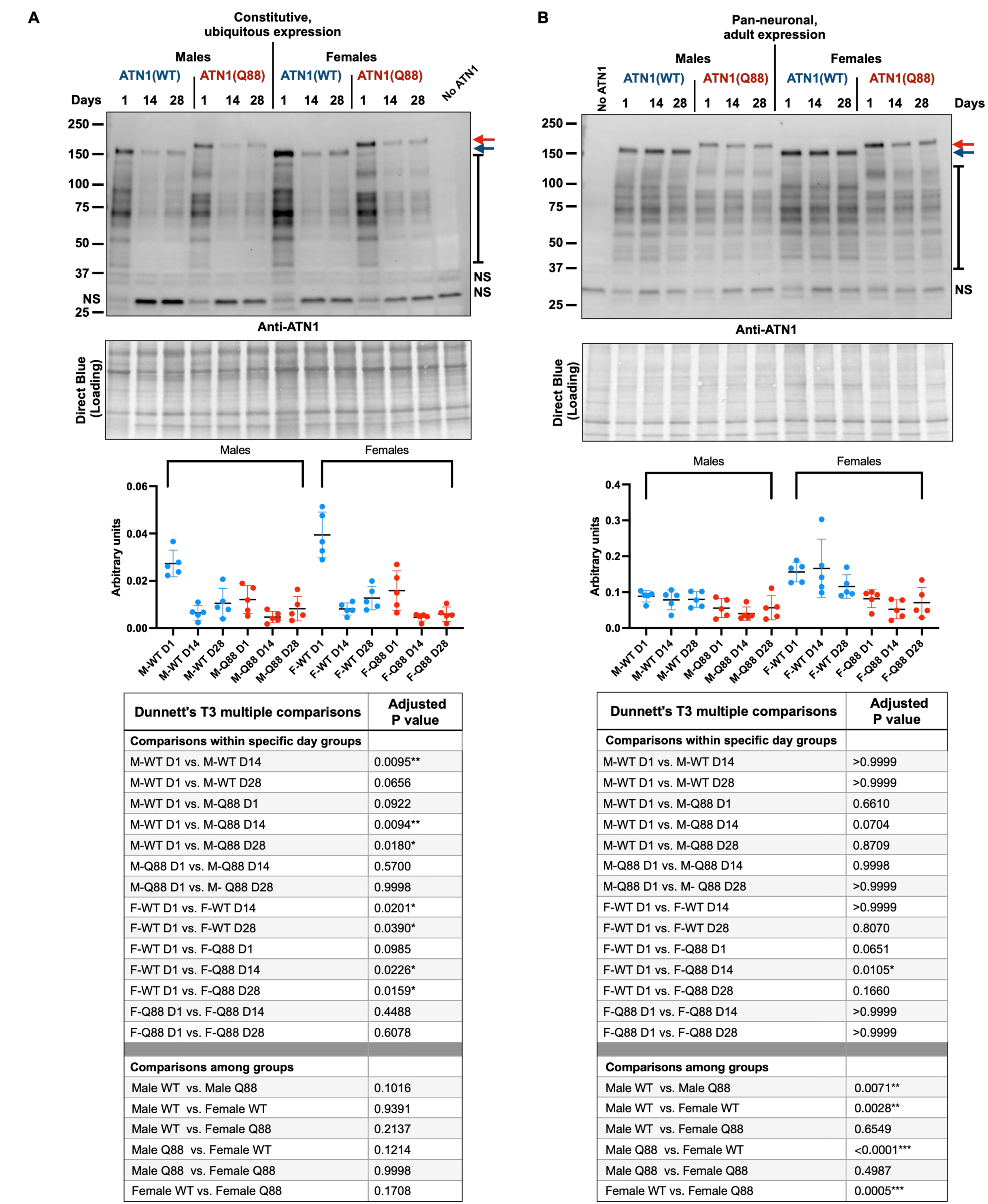
Ubiquitous and adult, pan-neuronal expression of ATN1 protein. Western blots from male and female flies (heads, 10 per sample) expressing the indicated transgenes (A) ubiquitously or (B) in adult neurons, and aged for 1, 14 and 28 days. Top panel: representative blots from each expression pattern. Red arrow depicts polyQ-expanded ATN1; and blue arrow depicts wild-type ATN1. Black brackets represent likely proteolytic fragments of ATN1. NS: non-specific signal. Bottom panel: graph, ATN1 protein levels from N=5 independent repeats for all lanes. Statistics: Brown-Forsythe and Welch ANOVA tests. Asterisks indicate significant differences.

Expression in adult neurons also led to readily detectable ATN1 proteins (figure 3B). The likely proteolytic fragments that we observed with ubiquitous expression are again present with pan-neuronal, adult expression. With this expression pattern, we noted lower levels of ATN1 protein in males compared to females, specifically with ATN1(WT) (figure 3B and the statistics associated with it). Overall, we conclude that ATN1 protein is readily detectable in these new fly lines; that both WT and pathogenic forms of this protein are aggregation prone; and that ATN1 exists as various proteinaceous forms in this organism.

### Effect of ATN1 on the fly transcriptome

We next turned our attention to unbiased assessments to assist us with additional downstream studies of DRPLA in the fruit fly. We conducted RNA-seq analyses from male and female flies expressing either WT or pathogenic ATN1 in all tissues throughout development and as adults (sqh-Gal4). Even though there is no evidence that DRPLA is sex-dependent^20,40^, we continued to separate male and female flies to investigate the possibility of sex-specific events in our model. We utilized adult flies that were newly eclosed from their pupal cases.

We identified sex as a major driver influencing differences among samples (supplemental figure 3). Overall changes to the transcriptome are summarized in the Venn diagrams in figure 4A (supplemental tables 1 and 2 list the intersects for each sex). Protein quality control-related proteins (Hsp68 and Hsp70Ba) and fibrinogen-like protein (CG5550) are shared among all groups regardless of sex (supplemental tables 1, 2). Differences were more pronounced in male flies, which, on average, showed 2-or higher fold differences among the same groups of comparisons as female flies (figure 4A).

**Figure 4:**
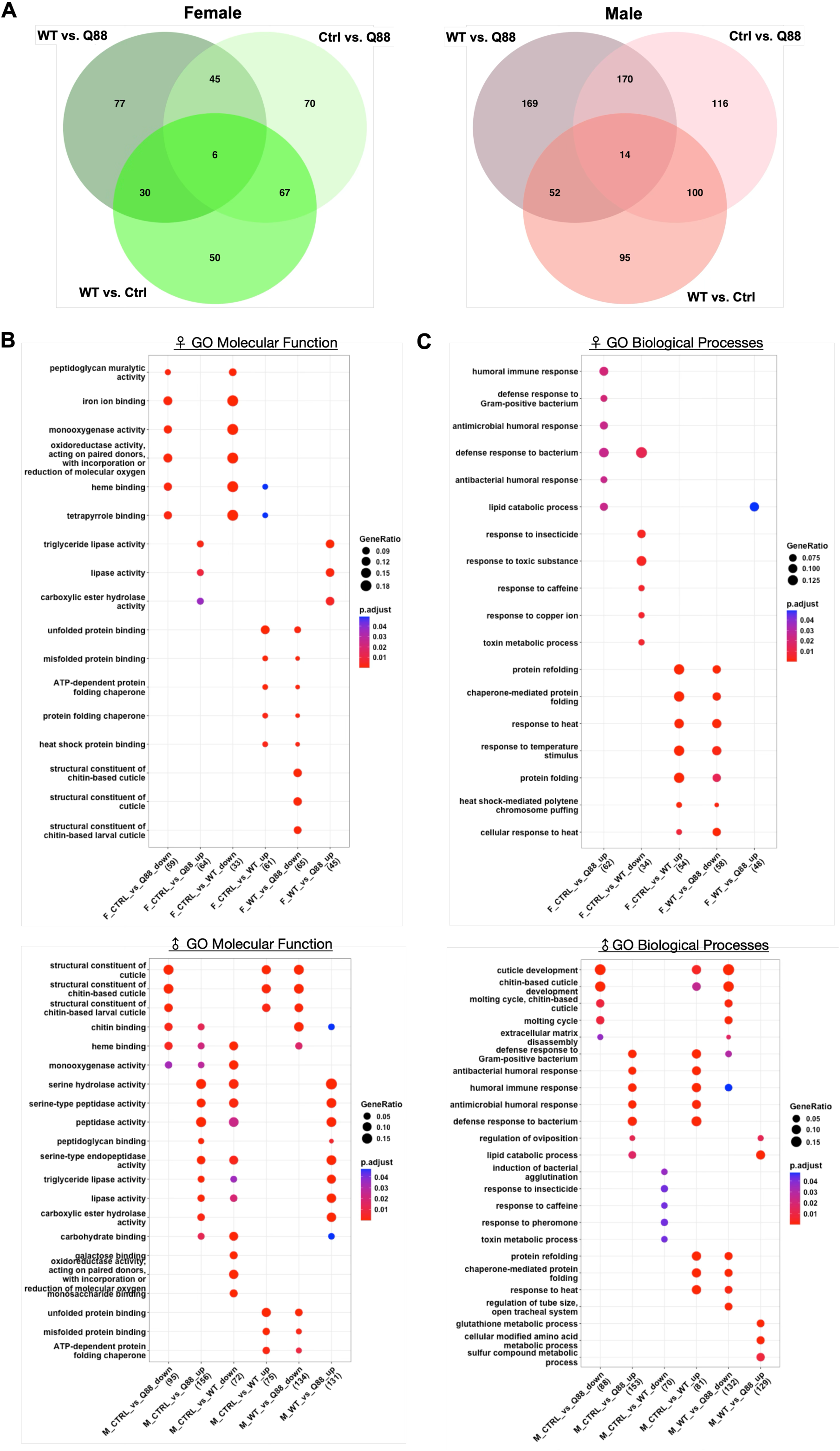
RNA-sequencing with constitutive, ubiquitous expression of ATN1 transgenes. A) Venn diagrams summarizing the number of differentially expressed genes between 1-day old female (left, green) or male (right, pink) flies expressing control, ATN1(WT) or ATN1(Q88) transgenes ubiquitously. B) Gene ontology (GO) analysis for molecular function of significantly up or downregulated genes in noted comparisons for female (top) and male (bottom) flies. C) GO analysis for biological processes of significantly up-or downregulated genes in noted comparisons for female (top) and male (bottom) flies. For both (B) and (C), the size of the circle for a given comparison correlates to the proportion of differentially expressed genes present in the comparison group relative to the total number of genes annotated to the pathway. The color denotes significance value, with blue being closer to 0.05 and red being closer to 0.01. N=5 independent samples per group analyzed. Expression directionality represented with the CTRL group as the reference.

When examining Gene Ontology (GO) enrichment of pathways involved in molecular functions (figure 4B) and biological processes (figure 4C), various pathways and processes emerged as significantly impacted by the expression of WT or pathogenic ATN1. In females, molecular functions such as peptidoglycan muralytic activity and tetrapyrrole binding were downregulated in ATN1(Q88) compared to Ctrl and in ATN1(WT) compared to Ctrl (figure 4B). Structural chitin-based functions were downregulated in ATN1(Q88) compared to WT, while showing no significant change relative to the Ctrl group (figure 4B). Male flies showed different responses in relation to females. These included downregulation of structural chitin-based functions in ATN1(Q88) compared to WT and Ctrl but upregulated in WT *versus* Ctrl flies (figure 4B).

Meanwhile, molecular functions such as heme binding, monooxygenase activity, triglyceride lipase activity, lipase activity and carboxylic ester hydrolase activity were upregulated in ATN1(Q88) flies compared to Ctrl in both male and female flies (figure 4B). Also shared between the sexes were unfolded protein binding, misfolded protein binding and ATP-dependent protein folding chaperone which were upregulated in ATN1(WT) compared to Ctrl but downregulated in ATN1(Q88) flies compared to ATN1(WT) (figure 4B).

We next analyzed the RNA-seq data to examine biological processes (figure 4C). Changes specific to ATN1(Q88)-expressing flies when compared to Ctrl were upregulation in processes involved in immune response (figure 4C). Shared processes included protein folding-related processes upregulated in female ATN1(WT) flies compared to Ctrl but downregulated in ATN1(Q88) compared to ATN1(WT) (figure 4C). Differences in biological processes were not wholly shared among female and male flies. For example, in males we again observed stronger responses in terms of structural processes such as cuticle development (figure 4C).

These data indicate specific responses by the fly environment to the expression of ATN1(WT) and ATN1(Q88) in male and female flies. Among the shared functions and processes were ones related to protein quality control and immune responses, which have been strongly implicated in the biology of disease of misfolded protein disorders and which have consistently been viewed as potential therapeutic targets^31,41–46^. We next focused on fly genes involved in protein quality control and immune response by excising them from RNA-seq datasets and determining their individual changes: might genes related to protein quality control and immune response be involved in this DRPLA model, providing new clues towards understanding its pathology and identifying potential therapeutic entry points? Figure 5 summarizes our findings.

**Figure 5:**
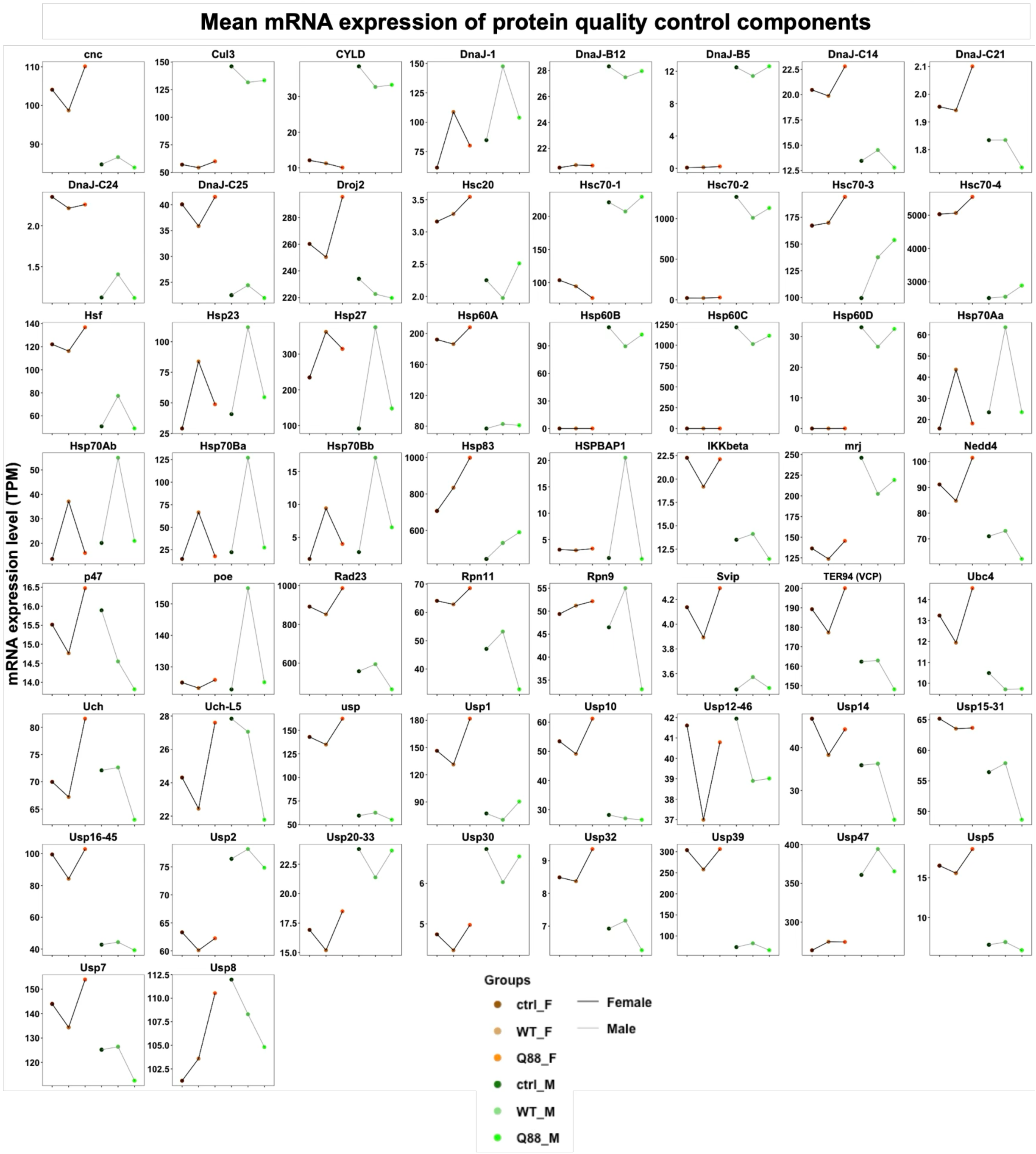
Transcriptome of protein quality control and immune response genes in DRPLA models. Line graphs of mean mRNA expression (Transcript *per* Million; TPM) of curated protein quality control genes across Ctrl, ATN1(WT) and ATN1(Q88) groups in male and female flies.

We found sex-specific differences for many of the protein quality control– and immune response-related genes that we assessed (figure 5). For some of these genes, the responses among groups in males and females were the opposite: e.g. with cnc (transcriptional regulator that controls the transcription of proteasome subunits, orthologue of mammalian NRF1 and NRF2), Hsf (activates transcription of heat shock proteins), a substantial portion of heat shock proteins (DnaJs, Hsps), Rad23 (a proteasome-binding protein that coordinates proteasome substrate delivery for degradation), and some deubiquitinases we noticed decreased levels when comparing control to ATN1(WT) samples and increased levels with ATN1(Q88) in females, while the opposite trend of response occurred in males. In a few other cases, we noticed similar trends between males and females with ATN1(WT) leading to higher levels than Ctrl and with ATN1(Q88) leading to levels closer to Ctrl (e.g., Hsp23, Hsp70). In yet other cases, females did not show differences in expression among different groups, whereas males did (e.g., CYLD, DnaJ-B12, HSPBAP1; Hsp60A showed nearly the opposite trend).

Altogether, these results suggest that the organism is responding to the presence of each human ATN1 protein in a distinct way that suggests physiological implications from each over-expressed transgene, but in ways that can help us distinguish the impact of the function of normal ATN1 compared to its disease-causing version in the future. Additionally, these transcriptomics behaviors, which clearly require additional future in-depth analyses, led us to examine how perturbing the levels of select genes from the ones shown in figure 5 may impact ATN1(Q88)-dependent pathology in flies. For these investigations, we turned to the fly eye.

### Pathogenic ATN1 causes eye anomalies in *Drosophila*

The *Drosophila* eye has become an invaluable tool for modeling neurodegeneration, especially to conduct large genetic screens^33–35,47^. We examined the impact of WT and pathogenic ATN1 on the external structure of the fly eye using a scale that we have reported before^5,32,47^ (figure 6A). Longitudinal examinations of the external structures of fly eyes expressing either variant of the ATN1 protein did not lead to observable defects at any time point evaluated (figure 6B; females are shown only, but males also did not show any anomalies through this assay). We confirmed that the intended protein was indeed expressed (figure 6B). Similar to the pattern of the ATN1 proteins in other tissues (figure 3), eye-restricted expression showed WT and pathogenic ATN1 migrating as expected, with a main band and multiple lower bands. The primary band diminished in intensity over time, likely also due to aggregation (figure 6B).

**Figure 6:**
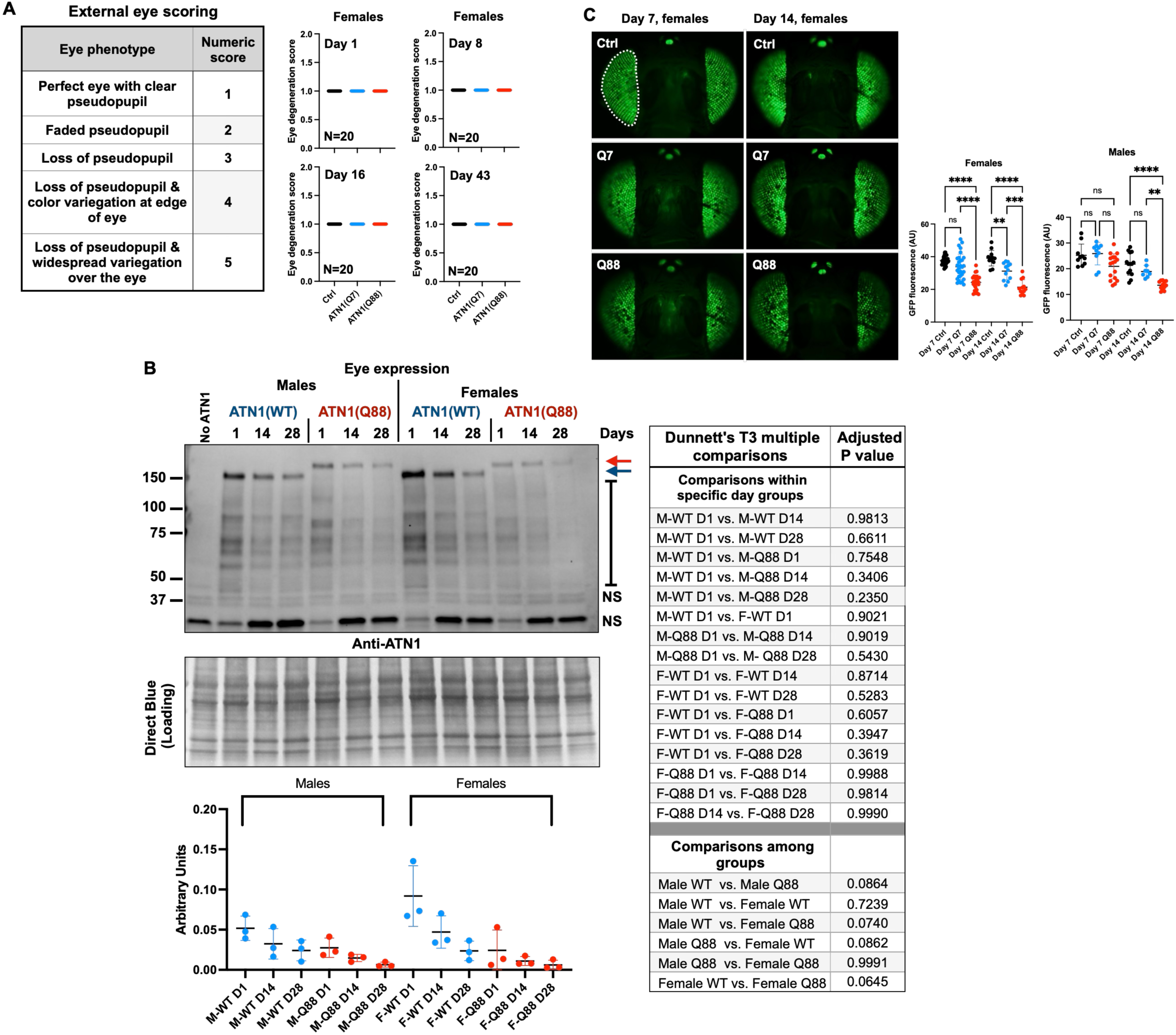
Effect of ATN1 protein expression in the *Drosophila* eye. A) Interpretation table for numerical scoring of eye phenotype (left), external eye scoring of female flies at day 1, 8, 16 and 43 (right). Control (Ctrl; black; GMR-Gal4 driver on the same background as ATN1 lines), ATN1(WT) (blue), ATN1(Q88) (red). N=20 flies per group. B) Western blots showing ATN1 expression levels in male and female flies following GMR-Gal4-dependent expression of the noted transgenes for 1, 14 or 28 days. N=3 independent repeats, 10 heads per sample. NS: non-specific signal. Statistics: Statistics: Brown-Forsythe and Welch ANOVA tests. C) CD8-GFP marker for assessing internal eye morphology in day 7 and day 14 female flies. White dotted outline shows area of quantification for GFP fluorescence, N ≥ 8 per condition. Statistics: Brown-Forsythe and Welch ANOVA tests. NS, not significant; **: p<0.005, ****: p<0.0001.

Next, we utilized a more sensitive assay to examine if ATN1 leads to any anomalies, potentially beneath the surface, again in fly eyes. We have reported and utilized this assay before; it centers on the presence of membrane-targeted GFP as a readout of internal eye structure. As ommatidia degenerate internally, GFP is lost, and the overall fluorescence of the eye diminishes^48^. As summarized in figure 6C, expression of pathogenic ATN1 leads to a significant loss of GFP fluorescence in fly eyes in both males and females by day 14. Expression of the WT version of ATN1 also has an effect in this assay, with significant toxicity on day 14 in females, but not in males. Based on these results, we conclude that pathogenic ATN1 is mildly toxic in fly eyes.

### Targeted screen for modifiers of DRPLA eye phenotypes

A major strength of fly-based models of disease is their utility towards screens to identify suppressors and enhancers of phenotype that can then be leveraged towards detailed understandings of both the biology of the disease and therapeutic opportunities for it in the clinic^49–56^. Thus, informed by results from our RNA-seq data we used the GFP-based assay described above to examine the effect of numerous protein folding-, immune– and proteasome-related genes on the previously observed ATN1(Q88) eye phenotype. To target these various genes, we used genetic mutations, over-expression and RNA-interference. Where possible, we employed multiple tools for each gene. These data are compiled in figure 7A.

**Figure 7:**
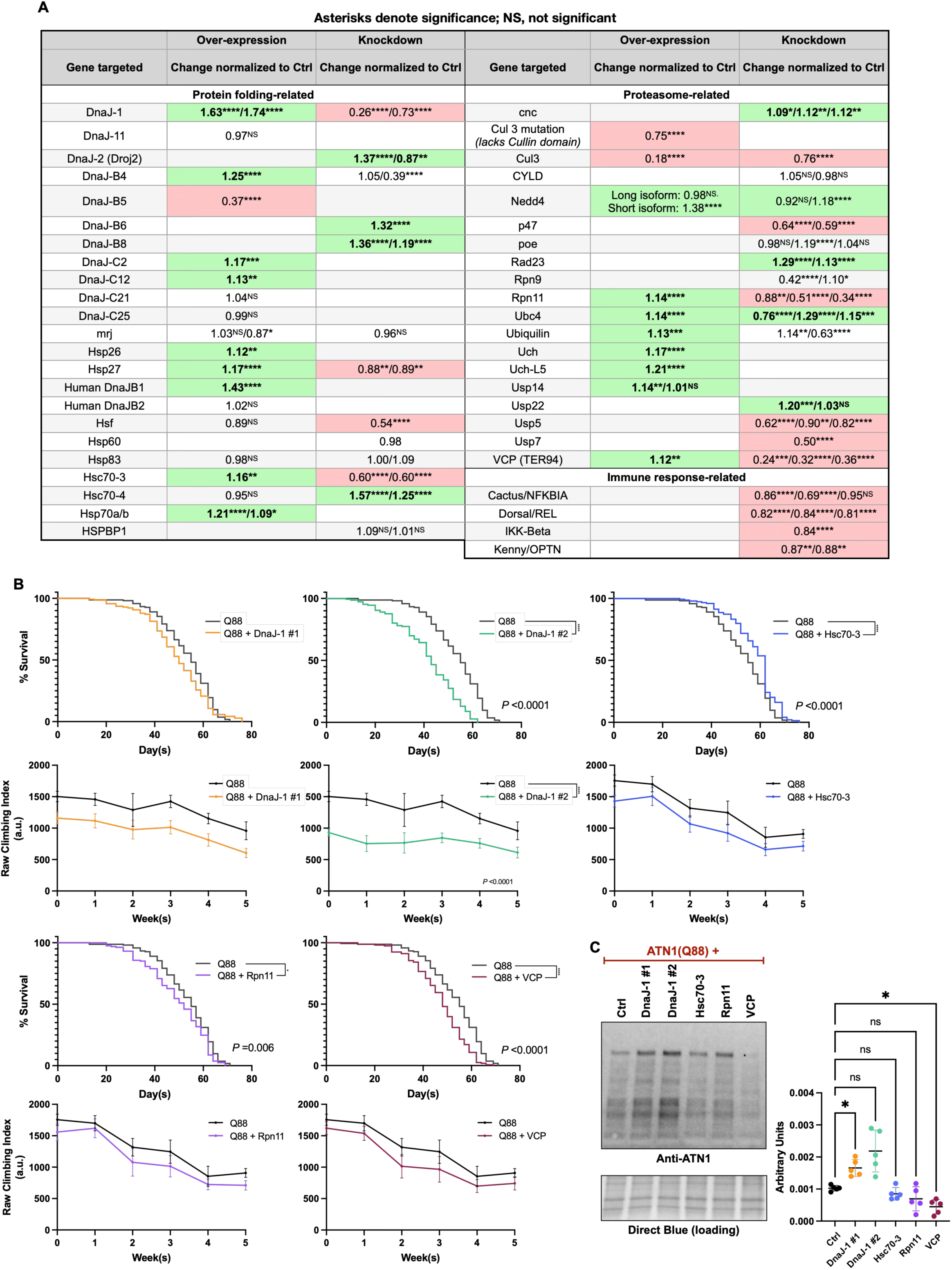
Genetic examination for modifiers of ATN1 toxicity. A) Summary of results of the genetic interventions using the CD8-GFP assay. N=20 per genotype. Statistics: Brown-Forsythe and Welch ANOVA. NS, not significant; *, p<0.05; **, p<0.01; ***, p<0.001; ****, p<0.0001. B) Longevity and motility data from the indicated genotypes. Statistics: Log-rank test, N≥100 (longevity). Simple linear regression, N=20 images per week, per genotype (motility). C) Western blots from the indicated genotypes, corresponding to panel (B). 10 heads per lysate from day 1 adults. Quantifications on the right are from data on the left and additional independent repeats. N=5. Statistics: Brown-Forsythe and Welch ANOVA tests. *: p=0.05.

Targeting of various chaperones, which assist with protein folding, had different outcomes. In some cases, they were helpful in reducing toxicity when overexpressed (e.g., DnaJ-1, DnaJ-B4, DnaJ-C2, DnaJ-C12, Hsp26, Hsp27, Hsc70-3, Hsp70). In other cases, overexpression had a negative impact (e.g., DnaJ-B5). And, in the rest of the cases tested they did not have a measurable impact. When targeted for knockdown by RNA-interference, perturbing some of these chaperones in fact led to a positive outcome (e.g., DnaJ-B6, DnaJ-B8, Droj2, Hsc70-4). We have observed differential effects from chaperones in the past in other polyQ diseases. Among them, Hsc70-4 was also problematic with the SCA3 protein, ataxin-3^31^.

Perhaps unsurprisingly, targeting the function of the proteasome (Rpn11, a component of the 19S cap of the proteasome that serves as the resident deubiquitinase and removes ubiquitin chains *en bloc* before substrate unfolding and degradation) led to worsening of the phenotype; increased levels of this same gene suppressed the phenotype. Perturbing some proteasome-related proteins such as VCP and p47 (which assist with substrate delivery to the proteasome) also enhanced the phenotype, whereas over-expression of VCP ameliorated it. Interestingly, RNAi of Rad23, another proteasome-associated protein that assists with substrate delivery, suppressed the phenotype. We have observed a similar effect in another polyQ disease, SCA3^33,34^, suggesting nodes of interaction among different polyQ diseases. Insofar as immune response-related genes were concerned, we observed uniform impact from their knockdown: in each case, the phenotype was worsened.

WB from some of the targets above hinted at correlates between changes in fluorescence and ATN1(Q88) levels when specific genes were targeted. On the one hand, knockdown of Hsc70-4, which led to reduced toxicity, appeared to occur while the levels of ATN1 were dramatically higher (supplemental figure 4A); worsening of toxicity from knockdown of VCP appeared to also occur alongside increased levels of ATN1. On the other hand, improvement of phenotype when Hsc70-3 was over-expressed was accompanied by somewhat reduced levels of ATN1(Q88). Similarly, over-expression of Hsp70a/b led to improvement concomitant with ATN1(Q88) protein level reduction. These data suggest that some of these protein quality control-related components have the capacity to impact the toxicity of ATN1(Q88) in relation with its protein levels (e.g., VCP, Hsc70-3, Hsp70a/b), whereas others in apparent indirect relation with it (e.g., Hsc70-4; supplemental figure 4A; supplemental figure 4B shows a network representation of the quality control components in figure 7 based on STRING database analysis).

Next, we ascertained the translatability of some of the findings from our eye-based screen to the whole fly. We over-expressed some of the strongest modifiers from figure 7A in all neurons, throughout development and in adults, and examined their longevity and motility. Figure 7B summarizes our findings from the over-expression of DnaJ-1 (we had access to 2 different over-expression lines for DnaJ-1 and used both), Hsc70-3, Rpn11 and VCP, each of which was protective in the eye model. Only the over-expression of Hsc70-3 had a modest, but significantly protective impact on the longevity of the DRPLA flies when co-expressed in all fly neurons. The other modifiers from the eye model either worsened longevity (DnaJ-1 overexpression line 2, Rpn11, VCP) or did not have a statistically significant effect (DnaJ-1 overexpression line 1). In terms of motility, over-expression of each selected gene led to worse motility from the very beginning. The one potential benefit we observed was from over-expression of DnaJ-1 line 2, which led to a slower decline in motility over time compared to the control. Lastly, we examined day 1 ATN1(Q88) protein levels from this expression pattern. As shown in figure 7C, DnaJ-1 led to higher levels of ATN1(Q88), Hsc70-3 and Rpn11 did not impact the levels of ATN1(Q88), whereas VCP was especially potent at reducing the levels of this protein in fly neurons.

The Gal4 driver that we utilized for the assays in figure 7B and 7C drove expression throughout development and during adulthood. Because of the significant mobility issues on day 1 with this driver when ATN1(Q88) was co-expressed with the lines that we selected (figure 7B, motility graphs), we wondered if developmental effects confound our conclusions. Therefore, we examined the most severe case, DnaJ-1, using adult-onset pan-neuronal expression (supplemental figure 5). This expression pattern lessened the reduced motility phenotype but did not change the overall outcome – neither longevity nor rate of decline in motility were improved with expression of these genes in whole flies (supplemental figure 5A), and ATN1(Q88) levels were somewhat elevated (supplemental figure 5B).

Lastly, due to the improvement in toxicity with over-expression of Hsc70-3, we also investigated its knockdown in whole flies. We initially sought to use the same Gal4 driver as in figure 7B, leading to constitutive pan-neuronal expression. However, Hsc70-3 knockdown in neurons throughout development led to pharate adult lethality with or without concurrent ATN1(Q88) expression (data not shown). Therefore, we used adult-onset pan-neuronal expression. Hsc70-3 knockdown significantly decreased both lifespan and motility in ATN1(Q88) flies (supplemental figure 6A and B). By week 3 of tracking motility, most flies in the knockdown group had died, preventing us from continuing the typical 5-weeks of observation. We did not observe statistically significant differences in ATN1(Q88) protein levels in these flies (supplemental figure 6C). Notably, Hsc70-3 knockdown also reduced lifespan in non-ATN1(Q88) expressing flies compared to wild-type controls; however, the mortality rate was significantly higher when ATN1(Q88) was co-expressed (supplemental figure 6D).

Collectively, we interpret these results to suggest that there are tissue-specific effects from protein quality control components on the toxicity of ATN1(Q88). In neurons, DnaJ-1 enhanced the toxicity of ATN1(Q88), likely by increasing the overall levels of the soluble DRPLA protein. The negative impact of Rpn11 and VCP does not appear to correlate with the overall levels of the offending protein. Hsc70-3 appears to be helpful in the longevity, albeit not in the motility, of DRPLA model flies, without a clear impact on the levels of ATN1(Q88) protein. The consistently helpful effect of this chaperone in eye and neuronal expression merits future attention to determine a mechanism of action.

### Endurance exercise has therapeutic potential in this DRPLA model

We recently reported on the utility of chronic endurance exercise in flies as a protective mechanism against some polyQ disorders (SCA2 and SCA6), but not others (SCA3)^30^. With the new *Drosophila* DRPLA models on hand, we wondered whether daily exercise is also protective in this disease. As additional justification, a curation of the RNA-seq data presented above revealed multiple changes in the levels of exercise-related genes in the presence of ATN1 transgenes including sestrin, multiple sirtuins, the recently characterized FNDC5 ortholog iditarod^57^, and various interactors of mTOR^58–61^ (supplemental figure 7). Here, we again observed sex-dependent differences (raptor, rictor, Sirt1, srl, tor, Tsc1), as well as changes that were specific to either ATN1(WT) or ATN1(Q88) (Idit, Mtor, raptor, REPTOR, rictor, Sirt7). Since some of these changes were in the direction opposite of typical exercise-induced expression (for example, the downregulation of foxo and idit in ATN1(Q88) males), we wondered if exercise may be beneficial at least in part by re-normalizing transcriptome alterations in our models.

We started by testing the well-established 3-week, ramped protocol that reproducibly leads to broad health span benefits in healthy flies, using the Power Tower^62,63^. Briefly, the Power Tower takes advantage of the involuntary negative geotaxis behavior in the fly to induce running for scheduled, increasing increments of time each week^63^. Both prior to and after completing training, the endurance (referred to as “runspan”) of the flies is tested to determine any impact genotype and/or exercise status might have on phenotype. As flies age, their endurance naturally declines, but exercise-trained flies reproducibly decline more slowly than their unexercised siblings when chronic exercise is beneficial for a given model^30,62,64^.

On day 5 (pre-training), we observed no differences between groups that would later be exercise-trained, or not (figure 8A; supplemental figure 8 shows the benefits of exercise to a generally accepted background control line, w^1118^, which we used for all of our work). With this protocol, we observed a strong trend toward improvement in exercised control flies containing the Gal4 driver without ATN1 without reaching statistical significance. We did not observe benefits to endurance in ATN1(WT)-expressing flies. ATN1(Q88) flies trended towards improved endurance without reaching statistical significance (figure 8B). According to WBs, exercise had no effect on ATN1 levels in either WT or Q88 flies, but Q88-expressing flies overall had lower levels of ATN1 than WT (figure 8D), similar to the trend we observed with day 1 adult males in figure 3.

**Figure 8:**
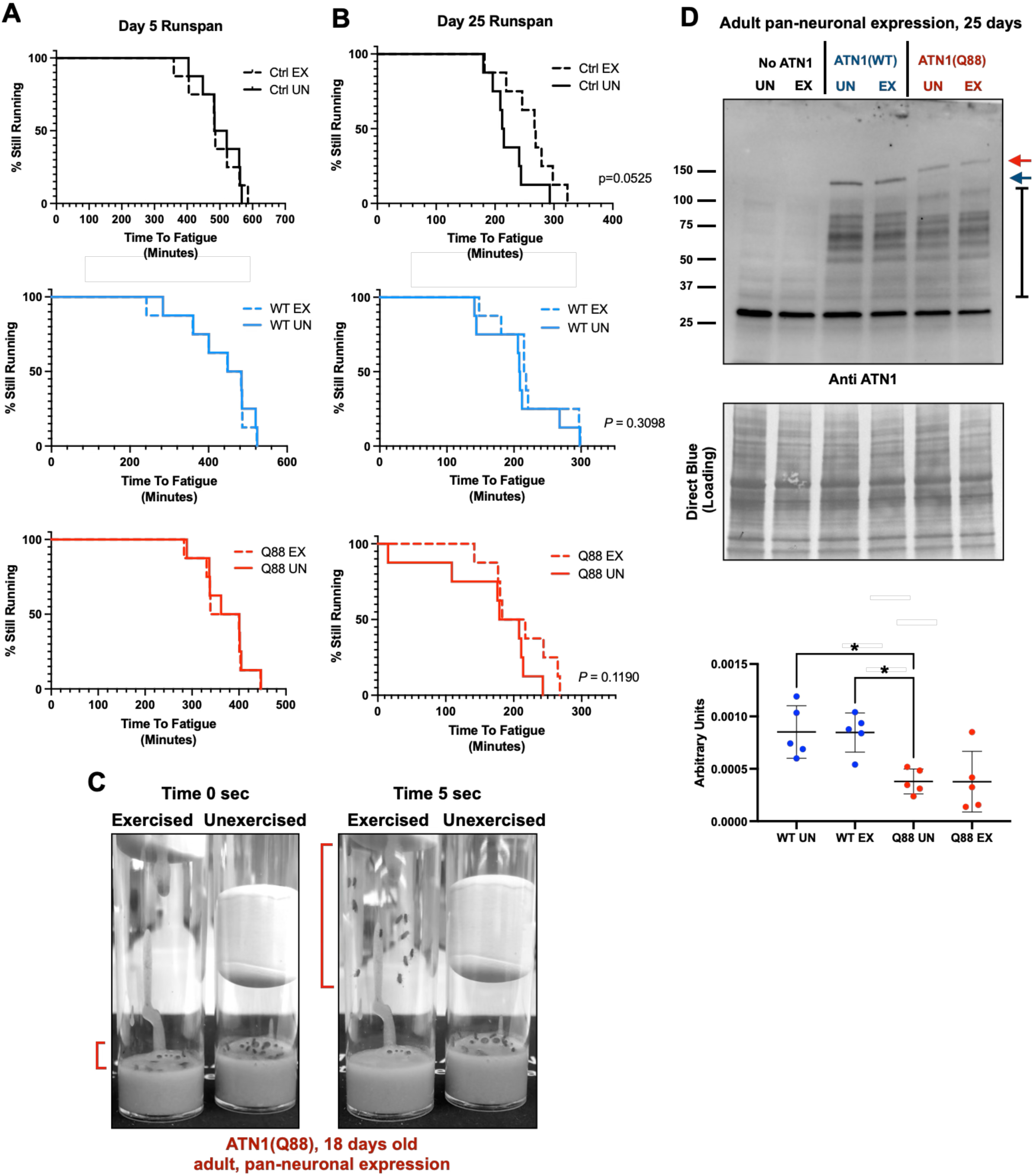
Endurance training in adult-expressing DRPLA models Day 5 (A; pre-training) and day 25 (B; post-training) endurance of male control (black), ATN1(WT) (blue) or ATN1(Q88) (red) flies. Dashed lines correspond to exercised flies, solid lines to unexercised flies. Statistics: Log-Rank tests, N=8 vials of 20 flies per genotype, per cohort. C) Snapshot of climbing ability in 18-day old ATN1(Q88) flies after completing 2-weeks of the chronic exercise protocol. Red brackets indicate height of exercise flies in their vials at start (left, 0 seconds) and 5-seconds after initiation of climbing (right). D) Western blot showing ATN1 levels post-training (day 25) under indicated conditions (top) and quantifications (bottom). Statistics: Brown-Forsythe and Welch ANOVA tests. N=5 independent repeats.

Interestingly, as we were assessing fly behavior with each passing week, we noticed that around the 2-week mark of training (18 days) the climbing ability of Q88 flies that were being exercised was visibly better than those that were unexercised (figure 8C). This observation, in addition to the trend toward improvement in >50% of the exercised Q88 vials in the post-training assessment (figure 8B) presented the possibility that a modified protocol, focusing around the 2-week mark, could be more beneficial in this model. Therefore, we next implemented an endurance exercise protocol with a 2-week duration. As with the 3-week training protocol, flies from each cohort were divided and tested pre-training for baseline endurance at 5 days-old. At this time point, Q88 cohorts showed reduced endurance compared to background controls (figure 9A). However, we again observed no differences between exercised and unexercised vials within groups pre-training (figure 9B). After 2 weeks of exercise, background controls and ATN1(WT) exercised flies trended toward increased endurance at day 18, but Q88 cohorts did not (figure 9C). Despite no benefit to endurance, Q88 exercised flies again had improved climbing ability compared to their unexercised siblings (figure 9D; a representative image from these data is shown in figure 9E). It is worth noting that when compared to non-ATN1 controls and ATN1(WT) flies, ones that expressed ATN1(Q88) and did not undergo training showed less reduction in endurance (compare day 25 runspans with solid lines in figure 9C), which may obscure a positive impact from exercise in this protocol. As previously observed, exercise did not significantly change the levels of ATN1 protein within cohorts, but Q88 flies again trended toward lower levels than WT expressing flies (figure 9F). Collectively, the data in figures 8 and 9 suggest that exercise is beneficial in DRPLA and that highly tailored exercise protocols are needed to harvest this potential.

**Figure 9:**
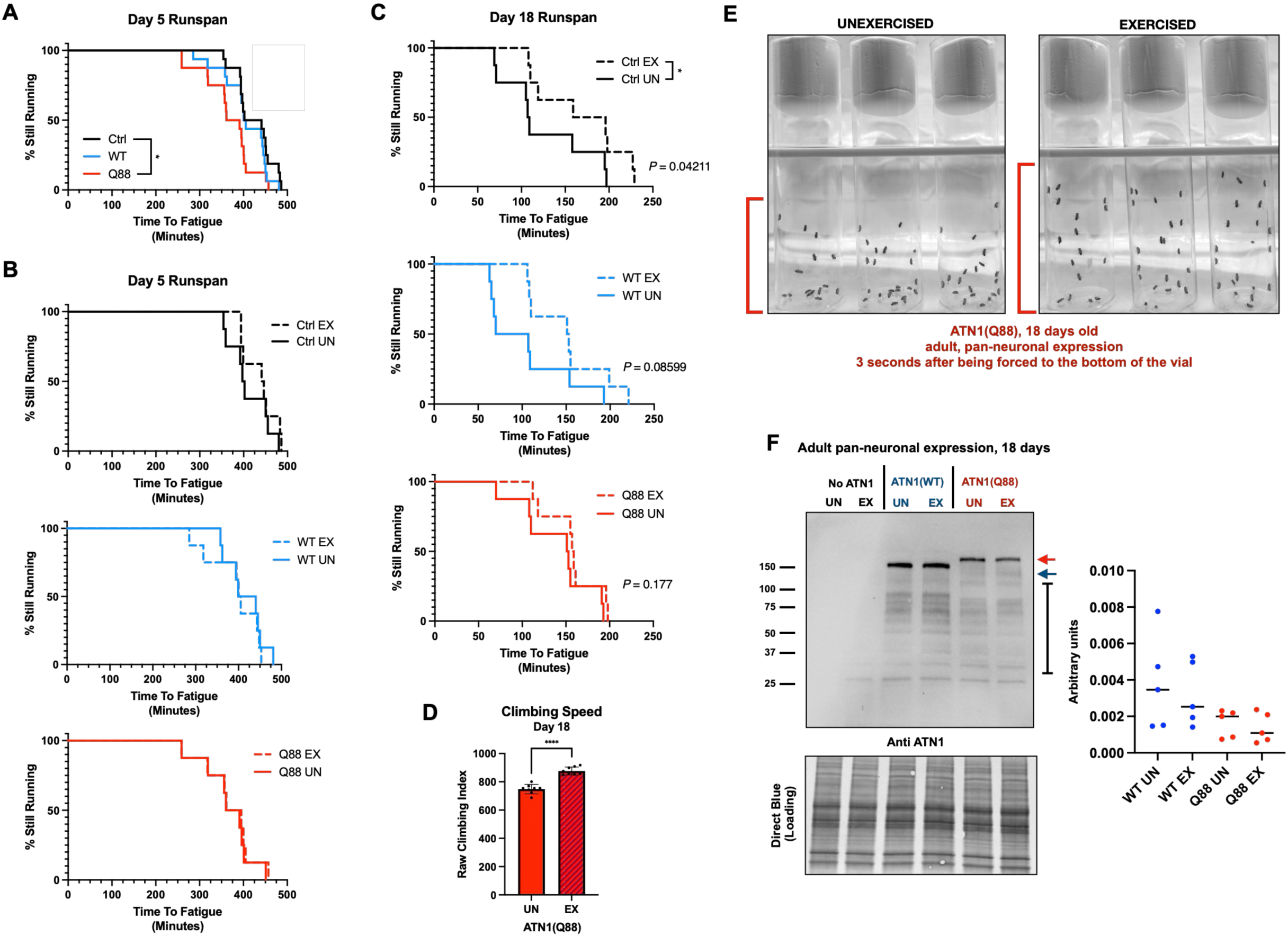
Modified endurance training in adult-expressing DRPLA models Combined day 5 (A), day 5 (B; pre-training) and day 18 (C; post-training) endurance of male control (black), ATN1(WT) (blue) or ATN1(Q88) (red) flies. Dashed lines correspond to exercised flies, solid lines to unexercised flies. Statistics: Log-Rank tests, N=8 vials of 20 flies per genotype, per cohort. *: p<0.05. D) Climbing ability in 18-day old exercised ATN1(Q88) flies. Statistics: Unpaired t-test. N=8 images per condition. ****: p<0.0001. E) Representative image from unexercised (left) and exercised (right) flies 3 seconds after initiation of climbing. Red brackets compare the maximum height of flies in the vials. F) Western blot showing ATN1 levels post-training (day 18) under indicated conditions (left) and quantifications (right). Statistics: Not significant by Brown-Forsythe and Welch ANOVA tests. N=5 independent repeats.

## DISCUSSION

We generated and conducted the initial characterization of novel transgenic *Drosophila* models of DRPLA, an incurable, age-related and inherited neurodegenerative disease that is under-investigated. Our new fly models show toxicity ranging from mild to severe in all tissues tested. A focused screen fueled by in-depth transcriptomics analyses identified protein quality control, proteasomal and immune response genes that act as modifiers of ATN1 toxicity in the fly.

Finally, exercise paradigms showed some therapeutic potential, reducing motility phenotypes caused by ATN1(Q88) expression in adult neurons. Collectively, we conclude that these new tools to understand DRPLA will prove beneficial to understand its biology and to develop therapeutics.

Few tools are available for studying DRPLA. Current models include cells^23^, transgenic mice^18^ and previously reported *Drosophila* lines^22,65^. Despite the beneficial insights that have come from these reagents, we lack effective treatment for DRPLA patients. Previous models have proven useful for exploring the pathogenic CAG expansions in the protein-coding region of the *ATN1* gene and our new models reflect features seen in the field: protein aggregation, motor defects, progressive degeneration and cellular toxicity. In addition to these characteristics, the findings that we reported reveal disruptions to protein homeostasis and many instances of altered gene expression in a sex-specific manner. Throughout our studies, we observed that both male and female flies were impacted by ATN1 over-expression; generally, males were more heavily impacted. While there is no clear evidence indicating sex-dependent effects in DRPLA patients^20,40^, our evidence of sex-dependent differences could inform future work to understand mechanisms that provide more protection to female flies compared to males expressing pathogenic ATN1 for the purposes of therapeutic intervention.

Protein quality control involvement is widely recognized across polyQ disorders^3,29,31,33,34,44,66^. The misfolding of polyQ-expanded proteins initiates a series of mechanisms that lead to cellular toxicity^1,3,44,67,68^. Aggregates of different sizes, especially oligomeric forms, can overwhelm and evade the cell’s degradative pathways, leading to significant cellular dysfunction^44^. While the consequences of this type of impairment vary across different diseases, several shared downstream effects have been reported. These include increased oxidative stress, compromised function of vital organelles such as mitochondria and the accumulation of polyQ aggregates alongside the dysregulation of the degradation of other proteins destined for elimination^1,3,52^. The changes in protein quality control transcript levels that we observed with the new DRPLA models (VCP, Rpn11, various chaperones) could indicate attempts by cells to overcome perturbations in protein quality control. We previously reported that VCP regulates the toxicity of another member of the polyQ family of diseases, SCA3^69,70^. In that case, VCP appeared to increase the toxicity of the SCA3 protein by enhancing its aggregation in various fly tissues^69,70^. Here, we found sex-specific differential expression of VCP (known as TER94 in the fly), which was markedly reduced in male flies expressing ATN1(Q88) (figure 5). In our DRPLA flies, over-expressing VCP reduced toxicity in fly eyes but enhanced it in neurons, concomitant with a reduction in ATN1 protein levels. VCP may well have distinct effects on the levels, aggregation and toxicity of the various polyQ proteins. VCP has many co-factors that regulate its function, activities and substrates, which may dictate the exact outcomes of its impact in a disease– and tissue-specific manner^71,72^. Its importance to quality control pathways can be appreciated by its central role among the genes we selected to interrogate genetically for this work, bridging folding-related proteins with the degradative machinery (e.g., supplemental figure 4B). The lineup of fly models of polyQ diseases that we have generated and reported^5,27–29,35^ (SCAs 3, 6, 7, 17, DRPLA and SBMA), as well as additional ones generated more recently (HD, SCAs 1, 2, as well as additional lines for SCA6 and SBMA), will serve as useful platforms for future work to dissect the role of VCP among all polyQ diseases and to understand its partnering options in each case.

The proteasomal subunit, Rpn11 is critical for protein degradation by removing ubiquitin chains from proteins anchored on the proteasome^73^. We observed sex-specific differential expression of Rpn11 (figure 5), which suggests that its expression and, potentially, its regulatory functions in protein degradation, differ by sex. A differential response could reveal unique insights into sex-specific vulnerabilities to proteostasis disruptions. This finding is particularly notable as our transcriptomic analysis revealed sex as a significant factor driving differences across samples (figure 4). Despite these sex-specific variations, however, the primary pathways of dysregulation appeared to be largely conserved (figure 5). Regarding the differential expression of proteasomal genes in our models, western blots for cyclin A – a cell cycle protein whose levels are tightly regulated via the ubiquitin-proteosome-system – revealed higher levels of cyclin A protein with the expression of ATN1(Q88) in neurons (supplemental figure 9). This suggests inefficiency of proteasomal activity in DRPLA, an additional finding the requires future investigation.

Another interesting set of proteins whose targeting has been reported to impact different polyQ proteins are DnaJ-1, Hsc70-3 and Hsc70-4. DnaJ-1 acted as a potent suppressor of SCA3 and SCA6 in *Drosophila*^29,35^. Here, we observed it to suppress toxicity in the fly eye but not in neurons, suggesting specific functions from this small co-chaperone in different polyQ diseases. SCA3 and SCA6 are significantly more toxic in the fly compared to DRPLA – ubiquitous expression of SCA3 and SCA6 with a polyQ repeat between 70-82 is developmentally lethal^29,31,35,69^, compared to this DRPLA model with Q88 repeats where adult flies emerge successfully and survive to ∼60 days (figure 1). Because DnaJ-1 is protective against high toxicity from SCA3 and SCA6 but not against DRPLA, perhaps it represents cellular processes that respond differently to misfolded proteins with varying toxicities.

Hsc70-4 was reported before to enhance the toxicity of SCA3^31^. Additionally, Hsc70-4 is upregulated as part of the cellular stress response in *Drosophila* models of Huntington’s disease, where it again was linked to enhanced toxicity^31,66^. We observed a similar inclination with DRPLA. In our DRPLA flies, Hsc70-4 expression was upregulated under ATN1 conditions (figure 5) and its knockdown ameliorated eye phenotype (figure 7A). suggesting that, similar to its effect in SCA3 flies, Hsc70-4 exacerbates DRPLA. Hsc70-3, a paralogue of Hsc70-4, was also upregulated in DRPLA flies (figure 5), yet its over-expression had positive impact on DRPLA in fly eyes and neurons, whereas its knockdown enhanced phenotypes (figure 7B and supplemental figure 6). We conclude that Hsc70 paralogs have differing roles in polyQ diseases and deserve future attention to map their functionalities in these disorders in a disease protein– and tissue-specific manner.

We noted mild toxicity from ATN1(WT) in some, but not all, of the tissues tested. Toxicity from ATN1(WT) was significantly milder than that of the disease-causing version; in some instances, we even observed improved behavior by flies expressing ATN1(WT). It is not uncommon for the normal versions of polyQ proteins to show some toxicity when they are over-expressed in the fly. Among polyQ diseases where this phenomenon has been reported are SCA7, SCA17, and SBMA^5,27,28^. This is likely due to the exogenous protein perturbing the function of other endogenous genes in the fly. The fly orthologue of ATN1 is Grunge, a nuclear repressor protein. The two proteins are not well conserved, with 24.3% similarity and 56.3% gaps (supplemental figure 10). Still, ATN1 could perturb the function of Grunge in tissues and processes where it is most important, especially during development^74,75^. We did not observe developmental issues from ATN1(WT) or (Q88). However, that does not mean that developmental issues that went undetected could not have had an impact later in the life of the fly. This point becomes particularly salient considering the possibility of developmental issues from pathogenic polyQ proteins^76–78^, which accentuates the need of developmental studies in our new model and in other polyQ diseases.

In other *Drosophila* models of polyQ disorders, chronic exercise was beneficial with varying degrees of benefit depending on the model being examined^30^. In DRPLA, we observed similar patterns; the benefits of exercise training seemed to be dependent on both the type of assessment being measured (chronic endurance or acute speed), as well as the duration of the training regimen. With DRPLA flies, a 2-week training period was sufficient to improve climbing ability in exercised ATN1(Q88) flies (figure 8C and 9D, E). Typically, 2 weeks of training is insufficient for healthy, wild-type flies to gain the full benefits of exercise using the Power Tower, which usually become apparent by day 21 of training^62^. Successful endurance training using the Power Tower relies heavily on an inducible running behavior (negative geotaxis) and the locomotor ability of the flies being trained. In flies with energetic deficits (e.g., muscle dysfunction, mitochondrial disease, or mutations that otherwise impair their ability to complete a standard protocol) the intensity of daily training could be titrated to meet the specific requirements and allowances of the disease model. We applied similar considerations to DRPLA flies by altering the duration of training (2 weeks *versus* 3 weeks) rather than intensity. Our observed trends toward improvement to various degrees depending on the duration of training, and on the energetic demands of the indices being measured for improvement (sustained endurance *versus* acute speed), suggest that training protocol adjustments are necessary for exercise to have a positive impact in DRPLA. This conclusion is relevant to studies in humans, where exercise has been suggested as a potential therapeutic option in the context of other polyQ disorders like Spinal and Bulbar Muscular Atrophy^79^. The intensity and duration of training play an important role in regular exercise being helpful or harmful – continuous, high-intensity exercise could be harmful in certain conditions or have little impact due to injury or excessive fatigue, whereas short intervals of high-intensity exercise can sometimes be beneficial^79^. In the DRPLA flies it is possible that, because endurance requires sustained activity rather than the short bursts required by climbing speed, flies struggle with meeting the demands of continuous running. As a result, they perform poorly on endurance assessments while still showing improvements in climbing ability. This same circumstance has been suggested in other fly models of disease^80^. Based on our collective data, chronic endurance exercise may be therapeutic in DRPLA with modified intensity and duration.

In conclusion, we described novel *Drosophila melanogaster* models of DRPLA that are posited to significantly enhance the understanding of the molecular and physiological mechanisms underlying toxicity and protection in this disease. Our collective assessments provide new insights into cellular responses to DRPLA and specify potential targets of intervention and exercise-related benefits. These new *Drosophila* lines can be used in conjunction with mammalian and cell models of DRPLA, facilitating progress in this disease. Additionally, they can be used alongside isogenic lines of other polyQ disorders in *Drosophila* to foster a comprehensive view of polyQ disease pathology and drive forward the search for effective treatments.

## METHODS

### Antibodies

The following antibodies were used: Anti-ATN1 (Santa Cruz, Cat. #sc-517594; 1:100 and 1:200), Anti-HA (Cell Signaling, Cat. #3724; 1-500 and 1:1000), Anti-Cyclin A (Developmental Studies Hybridoma Bank, ID#A12-c; 1:100), goat anti-Rabbit HRP conjugated secondary and goat anti-mouse HRP secondary (Jackson Laboratories, Cat. #115-035-144 and 115-035-146; 1:5000).

### Drosophila stocks

All flies used in the outlined experiments were housed at 25°C under controlled 50% humidity and a 12-hour light/dark cycle. They were fed a 10% yeast:sugar diet and flipped onto fresh food 3 times per week. In addition to the ATN1 models generated here, supplemental table 3 lists all the stocks used for crosses.

### Generating the new DRPLA models

We utilized the full-length sequence of human ATN1 with a CAGCAA doublet encoding the polyQ stretch. Genscript synthesized the cDNA with an in-line HA epitope tag at the 3’ end. The construct was cloned into pWalium10.moe and injected for insertion into site attP2. Following selection of transformants, their balancing and transfer into the w^1118^ background line, genomic DNA was extracted for PCRs to confirm insertion site and direction, and for sequencing to confirm line integrity and identity.

### Longevity

Survival was evaluated using cohorts of 100 age-matched flies for each tissue specific driver used. Flies were transferred to fresh vials 3 times per week, and deaths were recorded. In experiments selectively expressing ATN1 in adulthood, RU486 was mixed into the food during cooking at a concentration of 10mM and added at 20mL/L of food. Death events were graphed using GraphPad prism 9 and analyzed by log-rank tests. Any flies that escaped the vial or died from causes other than natural aging (e.g., crushed during flipping, etc.) were excluded from analysis.

### Motility

Motility was assessed by rapid iterative negative geotaxis (RING)^63^. Five vials containing 20 flies each (n=100 per cohort) were tapped down to the bottom of their vials to induce climbing via negative geotaxis response. Three seconds after initiation of climbing, a digital image was taken to capture the height of the flies in their vials. Longitudinal motility assessments were conducted at room temperature, with 20 images acquired weekly for 5 weeks. In experiments where ATN1 was expressed using the elavGS-Gal4 driver, flies were fed RU486 throughout the duration of the study. Raw image data were analyzed using ImageJ to calculate the average climbing speed (average height that flies were able to climb up their vials within 3 seconds) across a full week of images, and then averages were graphed using GraphPad prism software. Statistical analyses used are outlined in respective figure legends.

### Western blotting

For western blot analyses, 5 whole flies, or 10 fly heads (the exception was CD8-GFP crosses, where we used 20 fly heads per group) were homogenized in boiling lysis buffer (50mM Tris pH 6.8, 2% SDS, 10% glycerol, 100mM dithiothreitol (DTT)) using a tissue grinder (Axygen) designed for a 1.5 mL microfuge tube. Homogenates were then sonicated for 15s at 50% power, boiled for 13 minutes, and centrifuged for 7 minutes at 13,000 rpm at room temperature. The supernatant, excluding any fats, was loaded and electrophoresed through a pre-cast 4-20% Tris/Glycine gel (Bio-Rad), which was then transferred to a PVDF membrane (Bio-Rad). The membrane was then imaged using Chemidoc (Bio-Rad). Direct Blue was used to measure total protein levels. PVDF membranes were saturated for 10 minutes with 0.0008% Direct Blue 71 (Sigma-Aldrich) in 40% ethanol and 10% acetic acid, and then rinsed with a solution of 40% ethanol/10% acetic acid before being air dried overnight and imaged on Chemidoc (Bio-Rad).

Direct Blue signal was used as a loading control, unless stated otherwise. Quantifications of WB and Direct Blue images were conducted using non-saturated images in ImageLab (Bio-Rad) with global background subtraction.

### Soluble/pellet preparation

Five whole adult flies were homogenized in NETN lysis buffer (50mM Tris, pH 7.5, 150 mM NaCl, 0.5% Nonidet P-40) supplemented with protease inhibitor cocktail (S-8820, Millipore Sigma) and sonicated (50% power for 15 seconds on ice). Samples were then centrifuged at 20,000 x g at 4°C and the supernatant was transferred into a new microfuge tube on ice. The pellet was resuspended in 50 uL Laemmli buffer (12.5% 1M Tris, 20% Glycerol, 4% SDS, pH 7.5) and sonicated again at 50% AMP for 30 seconds. The resulting samples were then centrifuged for 2 minutes at 13,000 RPM and both soluble and pellet portions were supplemented with 6% SDS. Samples were then boiled 10 min, spun down briefly and loaded onto SDS-PAGE gels.

### Adult brain dissection and immunohistochemistry

Adult male and female flies were aged and dissected at timepoints corresponding with 0% (3 days on RU486), 25% (35 days on RU486), and 50% (50 days on RU486) mortality, as determined by survival curves with adult pan-neuronal expression of DRPLA. Brain dissection protocols were adapted from those outlined in Tito et al (2016)^81^. In summary, adult flies were anesthetized using light CO_2_ and briefly submerged in 100% ethanol to remove the wax coating on the outer cuticle before being transferred to a gel bottomed dissecting dish containing ice cold 1x Phosphate Buffered Saline (pH 7.2). Using sharp dissecting forceps, the head casing was removed to expose the brain tissue. Individual flies were then transferred to microfuge tubes containing 4% paraformaldehyde and fixed on rotation at room temperature for 60 minutes. Following fixation, the tissues were blocked for 2 hours (1xPBS + 0.3% Triton-X + 2% BSA) and incubated overnight in primary antibodies (Cell Signaling Technology, rabbit anti-HA, Catalog # 3724) on rotation at 4°C. Samples were then washed, incubated in secondary antibodies (ThermoFisher, goat anti-rabbit AlexaFluor 568, Catalog #: A-11011) and subsequently mounted in a 50% glycerol solution. Whole brain tissues were imaged using confocal microscopy at a magnification of 63X.

### RNA sequencing

RNA was harvested from 1 day old flies with TRIzol lysis buffer and isolated using the Invitrogen PureLink RNA Mini Kit with an ‘on the column’ DNAase digest. RNA quality was assessed on the Agilent 2200 TapeStation. Library preparation for sequencing was performed with PolyA selection. 150 bps paired-end libraries were sequenced on the Illumina NovaSeq. Prior to alignment the sequencing adaptors were trimmed using fastp v.0.23.1 and UMI-based de-duplication was performed using fastp v.0.23.1 simultaneously. Subsequently, trimmed and de-duplicated reads were then mapped to the *Drosophila melanogaster* BDGP6 reference genome available on ENSEMBL using the STAR aligner v.2.5.2b. Unique gene hit counts were calculated by using ‘featureCounts’ from the Subread package v.1.5.2. Differential gene expression was determined using the R package ‘edgeR’^82,83^ to calculate quantile-adjusted conditional likelihood (qCML), followed by an ‘exact’ test between two groups. Genes with an adjusted p-value < 0.05 and absolute log2 fold change > 1 were called as differentially expressed genes for each comparison. Gene Ontology analysis of differentially expressed genes was performed using ‘clusterProfiler’ version 4.6.2 in R. Genes were further annotated using ‘Drosophila_melanogaster_Ensembl_BDGP6’ from illumina and the ‘org.Dm.eg.db’ version 3.16.0^84^ package in R. Transcript per million counts were generated using sex normalized TMM counts. The entire dataset can be found as Supplemental dataset 1.

### CD8-GFP eye imaging and quantification

We conducted these assays following our previous publications^48,85^. In brief, flies containing GMR-Gal4 and UAS-CD8-GFP on the second chromosome and UAS-ATN1(Q88) on the third chromosome were crossed to selected RNAi, over-expression or mutation lines and their respective controls. Once adults eclosed from their pupal cases, flies were aged as indicated and heads were dissected and mounted onto microscope slides for 4X objective imaging on BX51 fluorescent Olympus microscope. Imaging settings were maintained constant across all imaging sessions and controls were imaged at the same time as experimental lines. Images were quantified using ImageJ free-hand selection tools in a blinded manner and mean brightness was recorded for each image.

### Endurance exercise training and runspan assessments

Ault male flies were collected within 48-hours of eclosing and flipped onto RU486 food for 3 additional days of aging before beginning endurance training on the Power Tower at 5 days old. Prior to the start of daily training, 16 total vials housing 20 flies each were tested for pre-training endurance, with 8 vials being arbitrarily labeled as unexercised and 8 as exercised. For this assessment, all vials were placed on the Power Tower with the vial plugs at the top of the vial. Negative Geotaxis was induced at regular 8 second intervals, and vials were removed from the Power Tower when 80% of the flies in a single vial had stopped responding to the induced negative geotaxis stimulus. The exact time a single vial was removed from the machine was noted, and the assessment continued until all vials had been removed. The time of removal for each individual vial was then plotted on a survival curve as a “runspan” to compare endurance between groups. The same assessment was done post-training at day 25. During training, flies were flipped onto fresh RU486 daily, and placed on the Power Tower for weekly, ramped time intervals starting at 2 hours per day in week 1, increasing to 2.5 hours per day in week 2, and ending with 3 hours per day in week 3. All vials were placed on the Power Tower to control for any non-specific effects caused by machine exposure, but the unexercised vials had their vial plugs pushed down low to prevent running. For the 2-week modified training protocol, all of the above steps were matched, but endurance training stopped after 2-weeks rather than continuing for 3.

### Statistics

Statistical analyses were conducted using GraphPad Prism. Specific statistical tests used throughout the paper are specified within the respective figure legends.

## AUTHOR CONTRIBUTIONS

MVP: conceptualization, data curation, software, formal analysis, funding acquisition, validation, investigation, visualization, methodology and writing and editing.

ON: conceptualization, data curation, software, formal analysis, funding acquisition, validation, investigation, visualization, methodology and writing and editing.

ROD: data curation, software, formal analysis, validation, investigation, visualization and methodology.

NCP: data curation, software, formal analysis, validation, investigation, visualization and methodology.

TK: data curation, formal analysis, supervision, validation, investigation and visualization. KL: data curation, formal analysis, validation, investigation and visualization.

AJB: data curation, formal analysis, supervision, validation, investigation,and visualization. WLT: data curation, formal analysis, supervision, validation, investigation and visualization. KR: conceptualization, resources, data curation, software, formal analysis, supervision, validation, investigation, visualization, methodology and writing and editing.

SVT: conceptualization, resources, data curation, software, formal analysis, supervision, funding acquisition, validation, investigation, visualization, methodology and writing and editing.

All authors contributed to the article and approved the submitted version.

