## Supplemental materials for "Insights into Dentatorubral-Pallidoluysian Atrophy from a new *Drosophila* model of disease"

### SUPPLEMENTAL MATERIAL LEGENDS

**Supplemental dataset 1:** Raw RNA-seq data.

**Supplemental table 1:** List of genes intersecting in female flies in figure 4A.

**Supplemental table 2:** List of genes intersecting in male flies in figure 4A.

**Supplemental table 3:** Fly stocks used in this study.

**Supplemental figure 1: Soluble/pellet fractionation of ATN1**
Shown are WB from two independent repeats of soluble/pellet fractionation as described in the "Methods" section. S: soluble; P: pellet; NS: non-specific staining. Flies were male.

**Supplemental figure 2: Expression of ATN1 in adult fly brain**
(Top) Representative image of female ATN1(Q88) whole-mount brain tissue following 25 days of RU486 feeding. 20X magnification. Yellow box: region magnified to observe ATN1 protein in brain tissue over time. (Bottom) Representative images of (left) female and (right) male control, ATN1(WT) and ATN1(Q88) whole-mount brain tissue at 63x magnification following 3, 35 and 50 days of RU486 feeding, N=5 observed per cohort. Yellow arrows indicate regions of punctate staining. All samples visualized using anti-HA antibody.

**Supplemental figure 3: Principal component analysis**
Principal component analysis (PCA) plot of Transcript *per* Million (TPM) normalized mRNA gene expression across control, ATN1(WT) and ATN1(Q88) groups in male and female flies. Proportion of variance is shown along the vertical and horizontal axes. The plot depicts sex-driven differences between samples accounting for 61% of variance.

**Supplemental figure 4: ATN1(Q88) protein levels from select modifiers of toxicity in the fly eye.**
A) WB using selected sets of male fly heads imaged for the data summarized in figure 7A. (Top) ATN1(Q88) expression levels with GMR expression of ATN1(Q88) alone (ctrl) or in the presence of the indicated transgenes. (Bottom) Total protein loading controls for each blot in top panel. 20 heads were lysed per sample. N=1 repeat. We used the same flies that were imaged for the

WBs in this panel, homogenized as a single group. B) STRING analysis of the genes targeted in figure 7. Line thickness indicates confidence, as noted in the legend. Active interaction courses are based on “Textmining”, “Experiments”, “Databases”, and “Co-Expression”. We selected “medium confidence: (0.400)”.

**Supplemental figure 5: Effect of adult only pan-neuronal expression of ATN1 and DnaJ-1 chaperone on longevity, motility and ATN1 protein levels.**
A) Longevity (top) and motility (bottom) with expression of ATN1(Q88) and DnaJ-1 overexpression line 1 (orange) or 2 (green) in adult nervous tissue. Statistics: Log-Rank tests. N≥100 per cohort. *, $p<0.05$; **, ****, $p<0.0001$. (longevity). Simple linear regression, N=20 images per week, per genotype (motility). B) Western blots from the indicated genotypes, corresponding to panel (A). 10 heads per lysate from day 1 adults. N=1.

**Supplemental figure 6: Effect of modifying Hsc70-3 expression in adult nervous system of ATN1(Q88) expressing flies.**
Longevity (A) and longitudinal motility (B) outcomes with expression of ATN1(Q88) (black), and ATN1(Q88) with Hsc70-3 over-expression (dark blue) or Hsc70-3 RNAi (light blue) in adult nervous tissue. N≥100 per cohort. Curve for Q88 and Q88 + Hsc70-3 show a smaller N due to flies still living at the time of figure generation (longevity). N=20 images per week, per genotype, simple linear regression (motility). C) Representative western blot from flies expressing ATN1(Q88) alongside the noted transgenes via RU486 feeding for 3 days (left). Quantifications (right) are from N=5 independent repeats. 10 heads per lane. Statistics: not significant by Brown-Forsythe and Welch ANOVA tests. The marked variability among samples comprising the Hsc70-3 RNAi group is likely due to: increased toxicity in these flies when Hsc70-3 is knocked down that may be perturbing various cellular pathways to different extents among different flies; and the fact that usually it takes longer than 3 days for the expression of proteins through the GeneSwitch system to equilibrate[79]. D) Longevity outcomes between Hsc70-3 RNAi lines with or without concurrent ANT1(Q88) expression. Statistics: Log-Rank tests. N≥100 per cohort. Curve for Hsc70-3 RNAi shows a smaller N due to flies still living at the time of figure generation.

**Supplemental figure 7: Expression levels of known exercise-related genes in DRPLA.**
Line graph of mean mRNA expression (Transcript *per* Million) of curated genes in exercise related pathways across control, ATN1(WT) and ATN1(Q88) groups in male and female flies.

**Supplemental figure 8: Wild-type exercise response to 3-weeks of endurance exercise training.**

Runspan comparisons between exercised (dashed lines) and unexercised (solid lines) from wild-type flies pre- (day 5) and post- (day 25) chronic exercise training. Statistics: Log-rank test, N=8 vials of 20 flies per condition.

**Supplemental figure 9: Insights into proteasomal function with adult nervous tissue expression of DRPLA.**

Western blot showing Cyclin A levels between non-expressing control and ATN1(Q88) expressing flies after 7 days of RU486 feeding. 10 heads per lane, N=2 for controls and N=3 for the DRPLA lane.

**Supplemental figure 10: Similarity between human ATN1 and fly ortholog.**

Sequence alignment of human ATN1 and the *Drosophila* ortholog “Grunge”.

Supplemental table 1: Female_intersects

| Ensembl | Gene | Entrez |
|---|---|---|
| **FBgn0001230** | Hsp68 | 42852 |
| **FBgn0013278** | Hsp70Ba | 44921 |
| **FBgn0034160** | CG5550 | 36883 |
| **FBgn0035412** | CG14957 | 38386 |
| **FBgn0051606** | CG31606 | 318842 |
| **FBgn0058469** | lncRNA:CR40469 | 5740321 |

Supplemental table 2: Male_intersects

| Ensembl | Gene | Entrez |
|---|---|---|
| **FBgn0001230** | Hsp68 | 42852 |
| **FBgn0001285** | Jon44E | 35853 |
| **FBgn0013278** | Hsp70Ba | 44921 |
| **FBgn0028519** | NA | NA |
| **FBgn0031034** | CG14205 | 32952 |
| **FBgn0034160** | CG5550 | 36883 |
| **FBgn0034407** | DptB | 37184 |
| **FBgn0039670** | CG7567 | 43479 |
| **FBgn0041581** | AttB | 36637 |
| **FBgn0052198** | CG32198 | 317909 |
| **FBgn0052865** | lncRNA:alphagamma-element:CR32865 | 3772597 |
| **FBgn0263597** | Acp98AB | 12798279 |
| **FBgn0265267** | CG18258 | 32708 |
| **FBgn0266949** | lncRNA:CR45400 | 19835792 |

| Supplemental table 3: Fly stock information | | | | |
|---|---|---|---|---|
| **Gene Symbol (Alphabetical)** | **CG Number** | **Stock Resource** | **Stock Number** | **Effect** |
| **cact** | CG5848 | BDSC | 31713 | Knockdown |
| **cact** | CG5848 | BDSC | 34775 | Knockdown |
| **cact** | CG5848 | BDSC | 37484 | Knockdown |
| **UAS-CD8-GFP** | Not Applicable | BDSC | 5137 | CD8-GFP expression |
| **Cnc** | CG43286 | BDSC | 25984 | Knockdown |
| **Cnc** | CG43286 | BDSC | 40854 | Knockdown |
| **Cnc** | CG43286 | BDSC | 32863 | Knockdown |
| **Cul3** | CG42616 | BDSC | 9936 | Overexpression |
| **Cul3 (mutation)** | CG42616 | BDSC | 9935 | Overexpression |
| **CYLD** | CG5603 | VDRC | GD15430 | Knockdown |
| **CYLD** | CG5603 | VDRC | KK101414 | Knockdown |
| **DNAJ-1** | CG10578 | BDSC | 32899 | Knockdown |
| **DNAJ-1** | CG10578 | BDSC | 32978 | Knockdown |
| **DNAJ-1 line 1** | CG10578 | BDSC | 30553 | Overexpression |
| **DNAJ-1 line 2** | CG10578 | FlyORF | F002258 | Overexpression |
| **DNAJ-11** | CG4164 | FlyORF | F003202 | Overexpression |
| **DNAJ-B1** | Not Applicable; Human DnaJ-B1 | BDSC | 53730 | Overexpression |
| **DNAJ-B1** | CG2887 | BDSC | 66983 | Knockdown |
| **DNAJ-B2** | Not Applicable; Human DnaJ-B1 | BDSC | 84936 | Overexpression |
| **DNAJ-B4** | CG7130 | FlyORF | F002684 | Overexpression |
| **DNAJ-B4** | CG5001 | VDRC | KK101532 | Knockdown |
| **DNAJ-B5** | CG2887 | FlyORF | F002411 | Overexpression |
| **DNAJ-B6** | CG8448 | VDRC | GD39126 | Knockdown |
| **DNAJ-B8** | CG7130 | VDRC | GD44395 | Knockdown |
| **DNAJ-B8** | CG7130 | VDRC | KK110526 | Knockdown |
| **DNAJ-C2** | CG10565 | FlyORF | F000235 | Overexpression |
| **DNAJ-C21** | CG2790 | FlyORF | F000438 | Overexpression |
| **DNAJ-C25** | CG7872 | FlyORF | F002616 | Overexpression |
| **dl** | CG6667 | BDSC | 27650 | Knockdown |
| **dl** | CG6667 | BDSC | 32934 | Knockdown |
| **dl** | CG6667 | BDSC | 34938 | Knockdown |
| **Droj2** | CG8863 | BDSC | 57382 | Knockdown |
| **Droj2** | CG8863 | BDSC | 36089 | Knockdown |
| **elav-Gal4** | Not Applicable | Daniel Eberl, University of Iowa | Not Applicable | Driver |

| | | | | |
|---|---|---|---|---|
| **elav-GS-Gal4** | Not Applicable | R.J. Wessells, Wayne State University | Not Applicable | Driver |
| **GMR-Gal4** | Not Applicable | BDSC | 8605 | Driver |
| **Hsc70-3** | CG4147 | FlyORF | F000956 | Overexpression |
| **Hsc70-3** | CG4147 | BDSC | 32402 | Knockdown |
| **Hsc70-4** | CG4264 | BDSC | 5846 | Overexpression |
| **Hsc70-4** | CG4264 | BDSC | 34836 | Knockdown |
| **Hsc70-4** | CG4264 | BDSC | 35684 | Knockdown |
| **Hsf** | CG5748 | FlyORF | F000699 | Overexpression |
| **HSF1** | CG5748 | VDRC | KK108851 | Knockdown |
| **Hsp26** | CG4183 | FlyORF | F000796 | Overexpression |
| **Hsp27** | CG4466 | FlyORF | F002489 | Overexpression |
| **Hsp27** | CG4466 | BDSC | 33007 | Knockdown |
| **Hsp27** | CG4466 | BDSC | 33922 | Knockdown |
| **Hsp60** | CG12101 | BDSC | 34729 | Knockdown |
| **Hsp60** | CG12101 | VDRC | GD18738 | Knockdown |
| **Hsp70Aa** | CG31366 | BDSC | 17624 | Overexpression |
| **Hsp70Ab** | CG18743 | BDSC | 15327 | Overexpression |
| **HSP83** | CG1242 | BDSC | 58469 | Overexpression |
| **Hsp83** | CG1242 | BDSC | 32996 | Knockdown |
| **Hsp83** | CG1242 | BDSC | 33947 | Knockdown |
| **HSPBP1** | CG10973 | VDRC | GD41463 | Knockdown |
| **HSPBP1** | CG10973 | VDRC | KK100869 | Knockdown |
| **IKKbeta** | CG4201 | BDSC | 35186 | Knockdown |
| **Jdp (DNAJ-C12)** | CG2239 | FlyORF | F002878 | Overexpression |
| **key** | CG16910 | BDSC | 35572 | Knockdown |
| **key** | CG16910 | BDSC | 57759 | Knockdown |
| **mrj** | CG8448 | BDSC | 30543 | Overexpression |
| **mrj** | CG8448 | BDSC | 30731 | Overexpression |
| **mrj** | CG8448 | BDSC | 66921 | Knockdown |
| **Nedd4** | CG42279 | BDSC | 31687 | Knockdown |
| **Nedd4** | CG42279 | BDSC | 34741 | Knockdown |
| **Nedd4 (long)** | CG42279 | BDSC | 81604 | Overexpression |
| **Nedd4 (short)** | CG42279 | BDSC | 81607 | Overexpression |
| **p47** | CG11139 | VDRC | KK107148 | Knockdown |
| **p47** | CG11139 | VDRC | GD17529 | Knockdown |
| **poe** | CG14472 | BDSC | 32945 | Knockdown |
| **poe** | CG14472 | VDRC | GD17648 | Knockdown |

| | | | | |
|---|---|---|---|---|
| **poe** | CG14472 | VDRC | KK108296 | Knockdown |
| **Prosalpha2** | CG5266 | BDSC | 44560 | Knockdown |
| **Prosalpha3** | CG9327 | BDSC | 55217 | Knockdown |
| **Prosalpha5** | CG10938 | BDSC | 34786 | Knockdown |
| **Prosalpha6** | CG4904 | BDSC | 34811 | Knockdown |
| **Prosbeta2** | CG3329 | BDSC | 67363 | Knockdown |
| **Prosbeta5** | CG12323 | BDSC | 34810 | Knockdown |
| **Rad23** | CG1836 | VDRC | GD30497 | Knockdown |
| **Rad23** | CG1836 | BDSC | 44031 | Knockdown |
| **Repo-Gal4** | Not Applicable | Vanessa Auld, University of British Columbia | Not Applicable | Driver |
| **Rpn11** | CG18174 | FlyORF | F000743 | Overexpression |
| **Rpn11** | CG18174 | VDRC | GD19272 | Knockdown |
| **Rpn11** | CG18174 | VDRC | GD19273 | Knockdown |
| **Rpn11** | CG18174 | BDSC | 33662 | Knockdown |
| **Rpn9** | CG10230 | BDSC | 34034 | Knockdown |
| **Rpn9** | CG10230 | BDSC | 67915 | Knockdown |
| **sqh-Gal4** | Not Applicable | Daniel Keihart, Duke University | Not Applicable | Driver |
| **TER94 (VCP)** | CG2331 | FlyORF | F001765 | Overexpression |
| **TER94 (VCP)** | CG2331 | BDSC | 31968 | Knockdown |
| **TER94 (VCP)** | CG2331 | BDSC | 32869 | Knockdown |
| **TER94 (VCP)** | CG2331 | BDSC | 35608 | Knockdown |
| **Ubc4** | CG8284 | BDSC | 15447 | Overexpression |
| **Ubc4** | CG8284 | VDRC | KK106600 | Knockdown |
| **Ubc4** | CG8284 | VDRC | GD35873 | Knockdown |
| **Ubc4** | CG8284 | BDSC | 64035 | Knockdown |
| **Ubqn** | CG14224 | BDSC | 83361 | Overexpression |
| **Ubqn** | CG14224 | BDSC | 33989 | Knockdown |
| **Ubqn** | CG14224 | BDSC | 33917 | Knockdown |
| **Uch** | CG4265 | FlyORF | F002766 | Overexpression |
| **UchL5** | CG3431 | FlyORF | F001113 | Overexpression |
| **Usp14** | CG5384 | FlyORF | F001032 | Overexpression |
| **Usp14** | CG5384 | FlyORF | F001366 | Overexpression |
| **Usp22** | CG4166 | BDSC | 28725 | Knockdown |
| **Usp22** | CG4166 | BDSC | 28725 | Knockdown |
| **Usp5** | CG12082 | VDRC | GD17567 | Knockdown |
| **Usp5** | CG12082 | VDRC | GD17658 | Knockdown |
| **Usp5** | CG12082 | BDSC | 31886 | Knockdown |

| | | | | |
|---|---|---|---|---|
| **Usp7** | CG1490 | BDSC | 34708 | Knockdown |
| **Stock Resource Abbreviates and Website Addresses** | BDSC: Bloomington *Drosophila* Stock Center - https://bdsc.indiana.edu FlyORF: Zurich ORFeome Project - https://flyorf.ch/ VDRC: Vienna *Drosophila* Resource Center - https://www.viennabiocenter.org/ | | | |

Supplemental figure 1

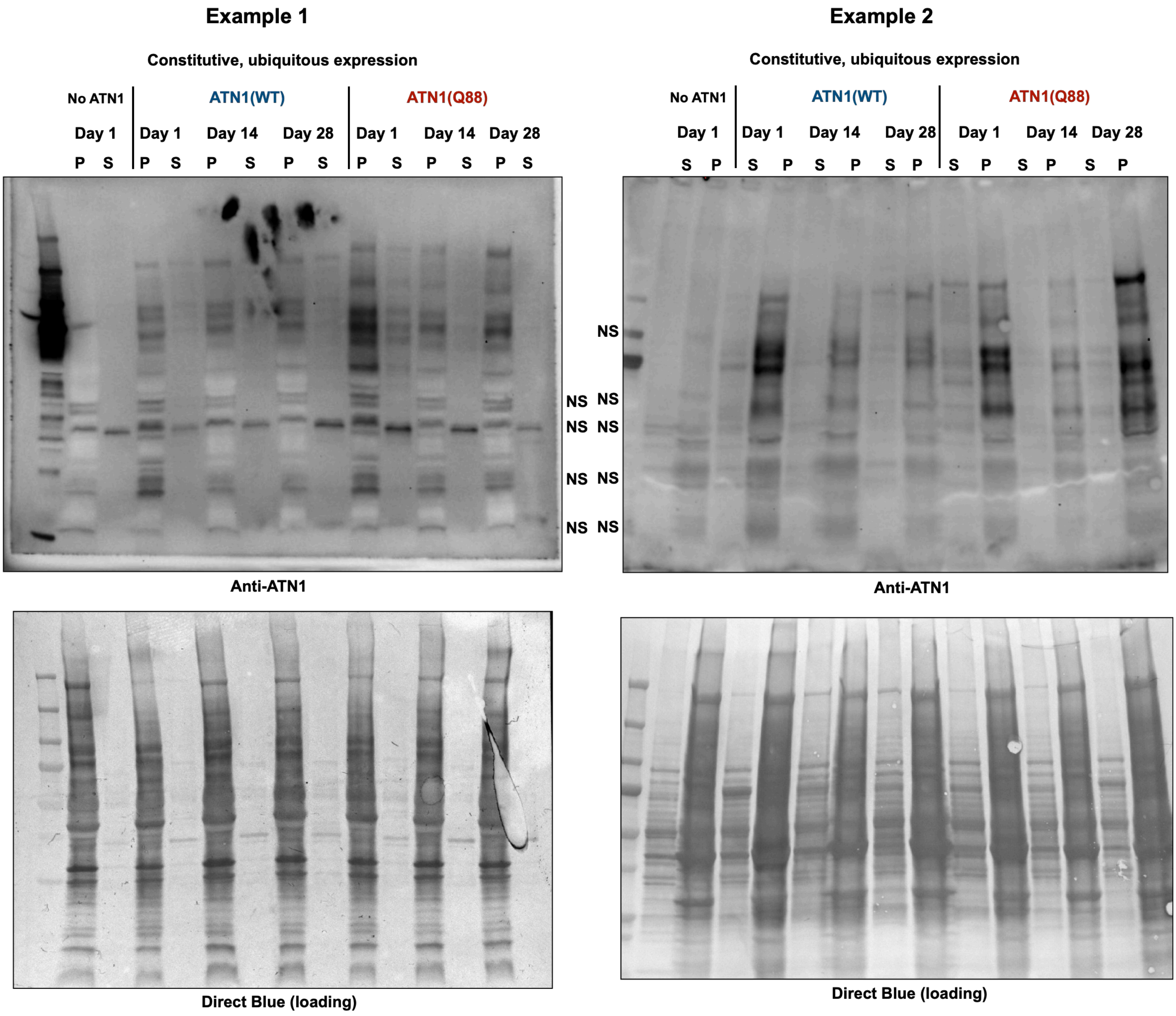

Supplemental figure 2

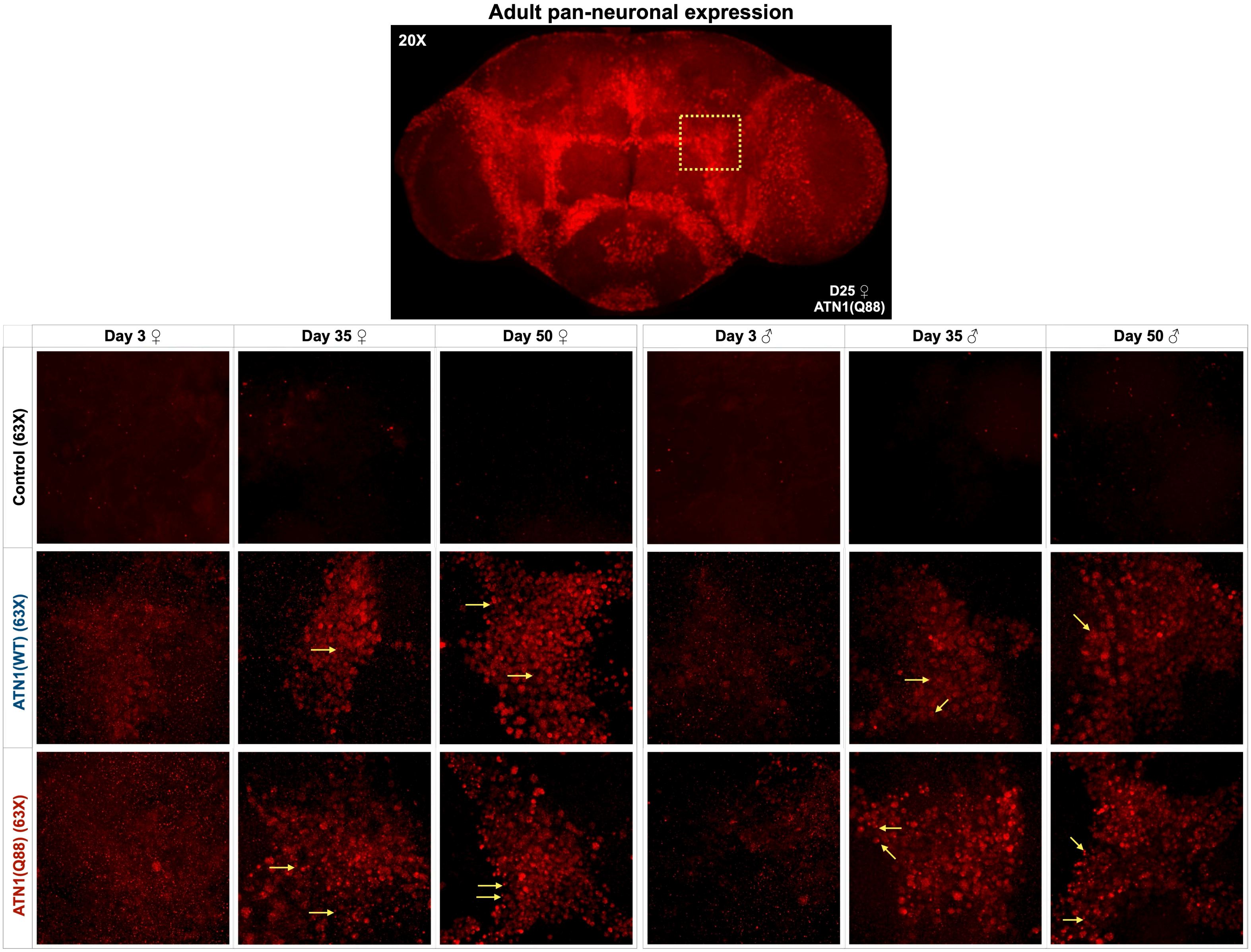

**Supplemental figure 3**

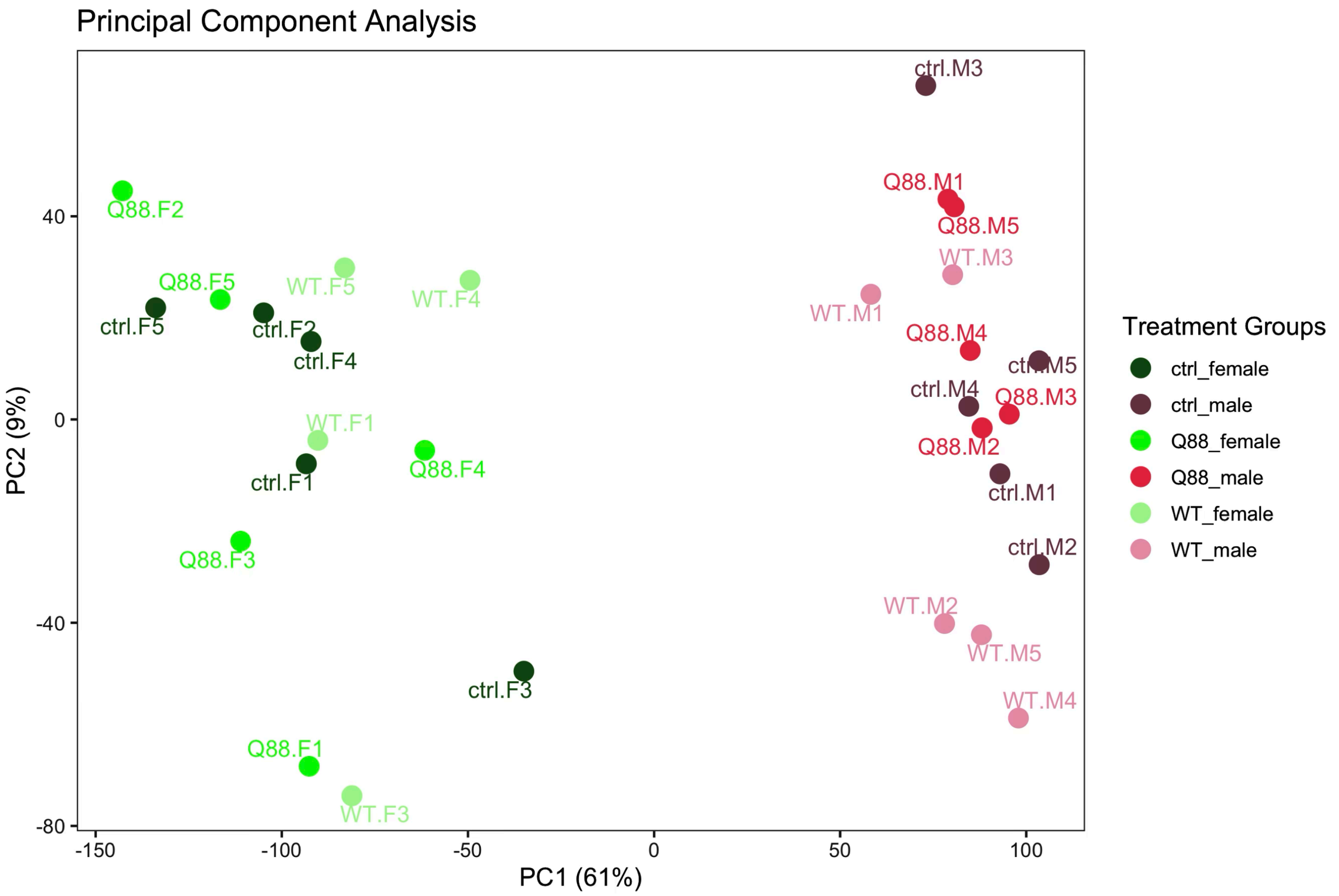

**Supplemental figure 4**

**A**

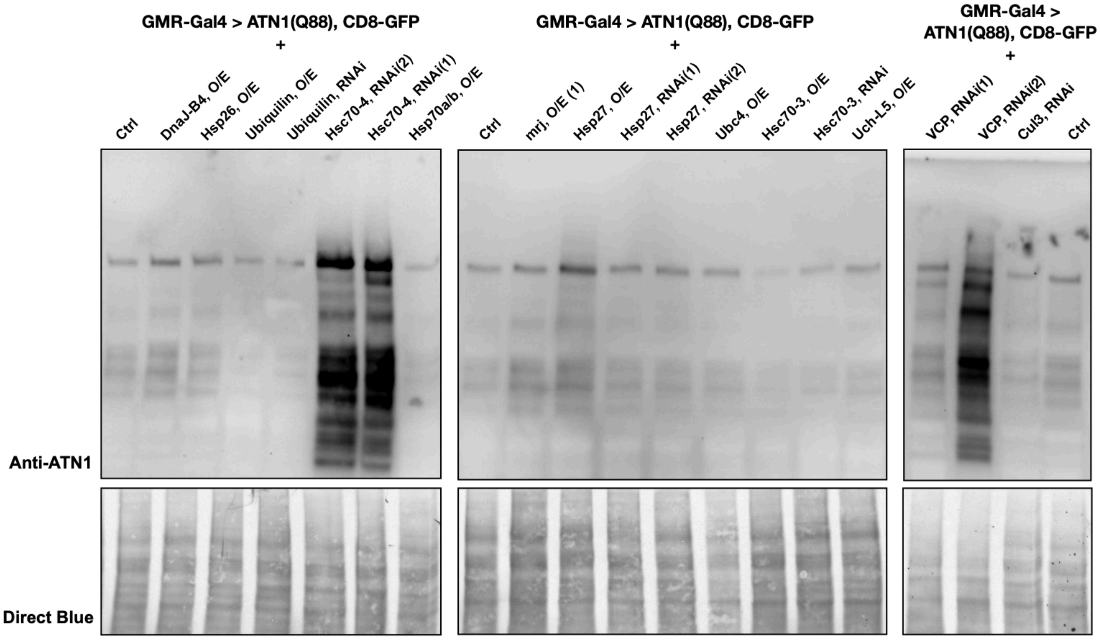


**B**

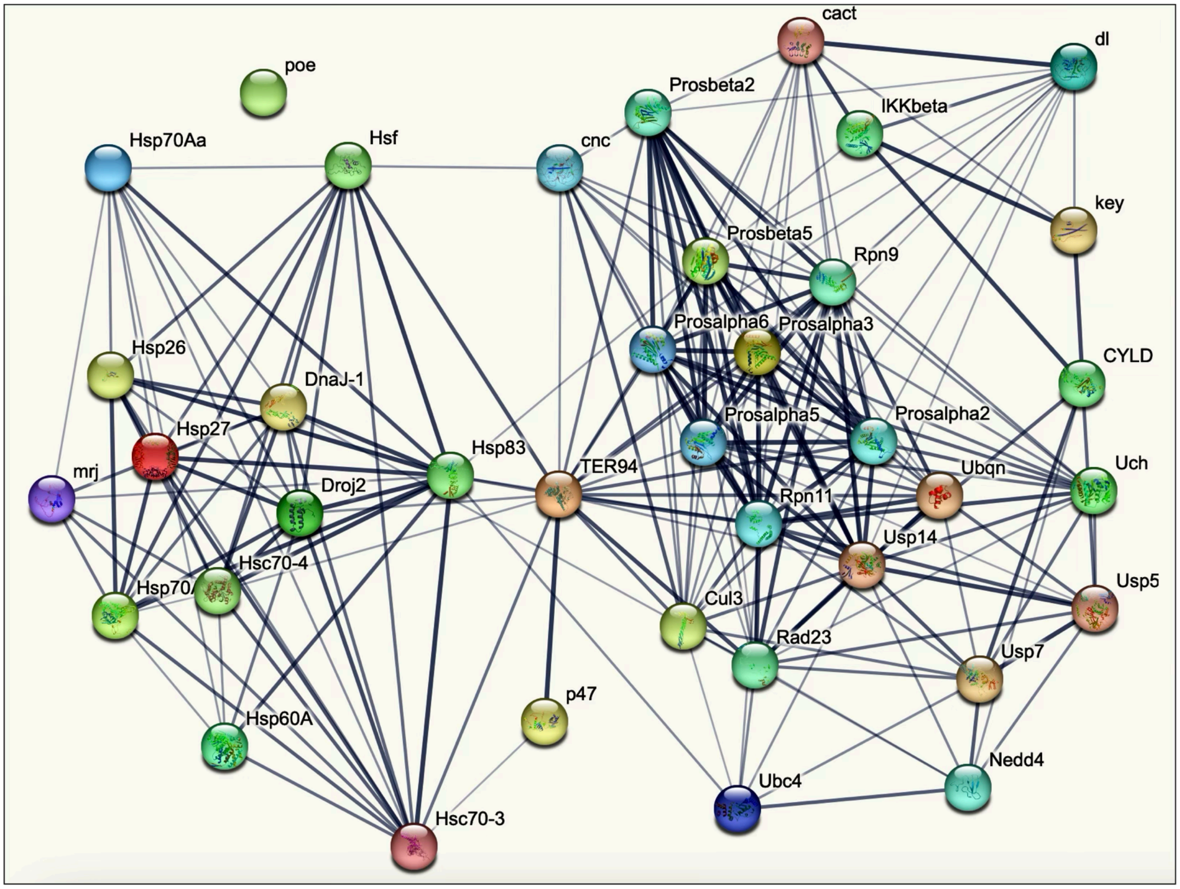


**Nodes:**

| Network nodes represent proteins | Node Color | Node Content |
| --- | --- | --- |
| *splice isoforms or post-translational modifications are collapsed, i.e. each node represents all the proteins produced by a single, protein-coding gene locus.* | *colored nodes:* *query proteins and first shell of interactors*<br>*white nodes:* *second shell of interactors* | *empty nodes:* *proteins of unknown 3D structure*<br>*filled nodes:* *a 3D structure is known or predicted* |

**Edges:**

| Edges represent protein-protein associations | Edge Confidence |
| --- | --- |
| *associations are meant to be specific and meaningful, i.e. proteins jointly contribute to a shared function; this does not necessarily mean they are physically binding to each other.* | *low (0.150)* *high (0.700)*<br>*medium (0.400)* *highest (0.900)* |

### Supplemental figure 5

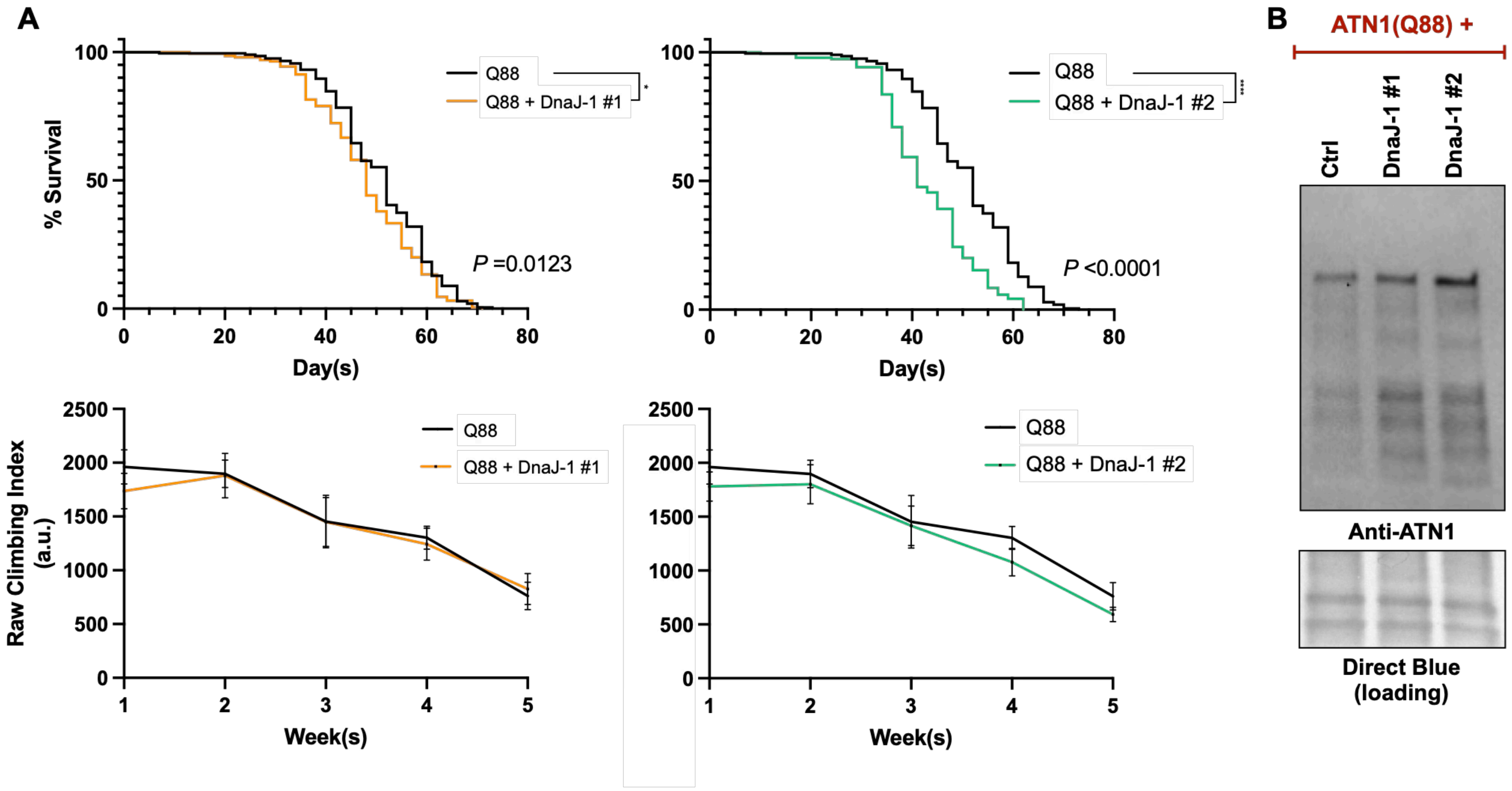

### Supplemental figure 6

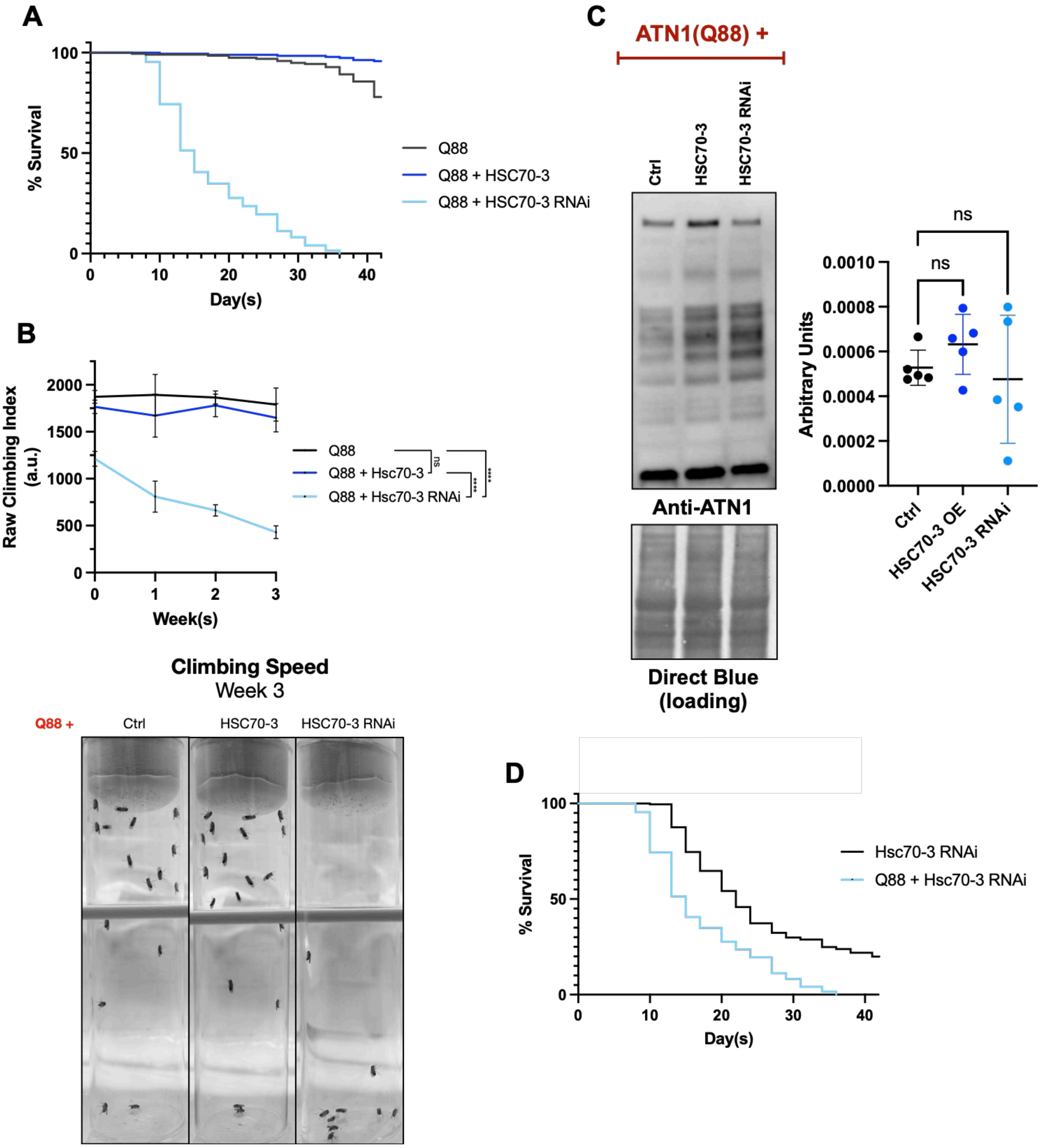

**Supplemental figure 7**

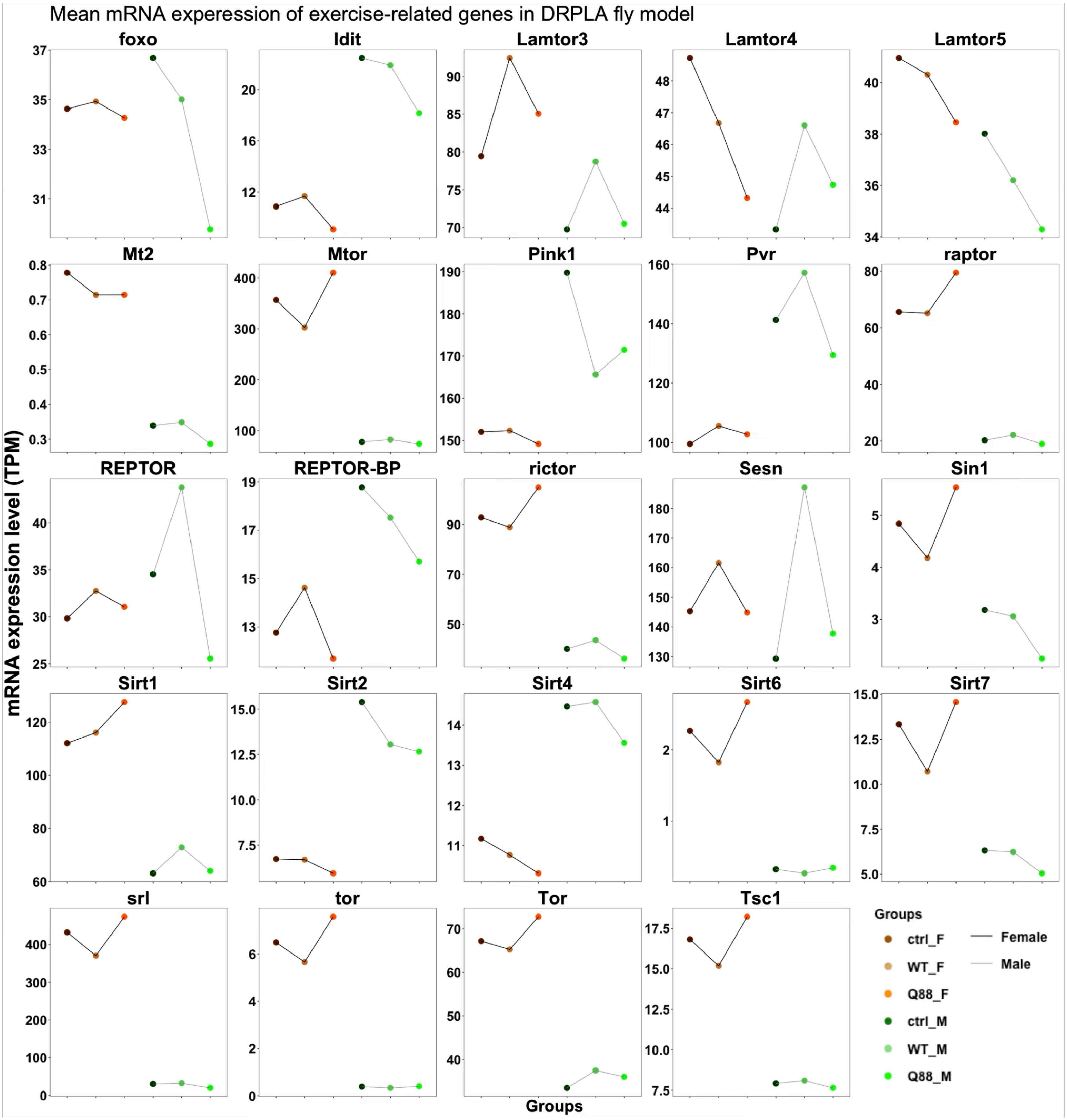

### Supplemental figure 8

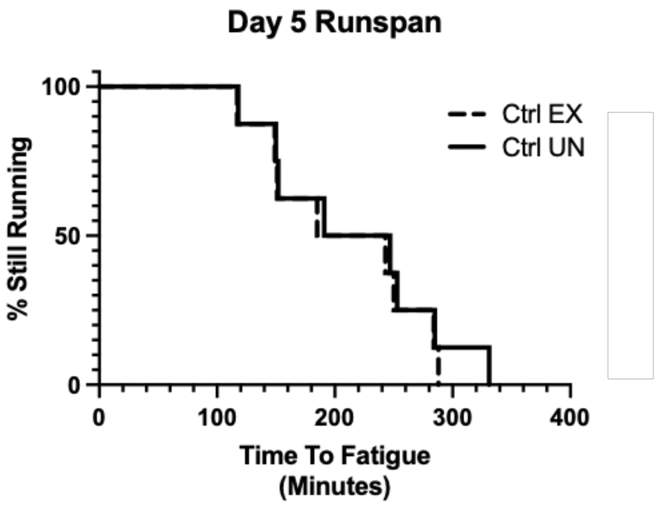


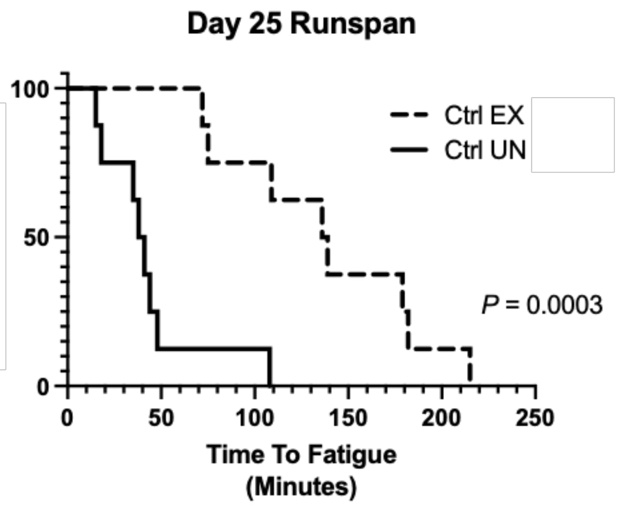

Supplemental figure 9

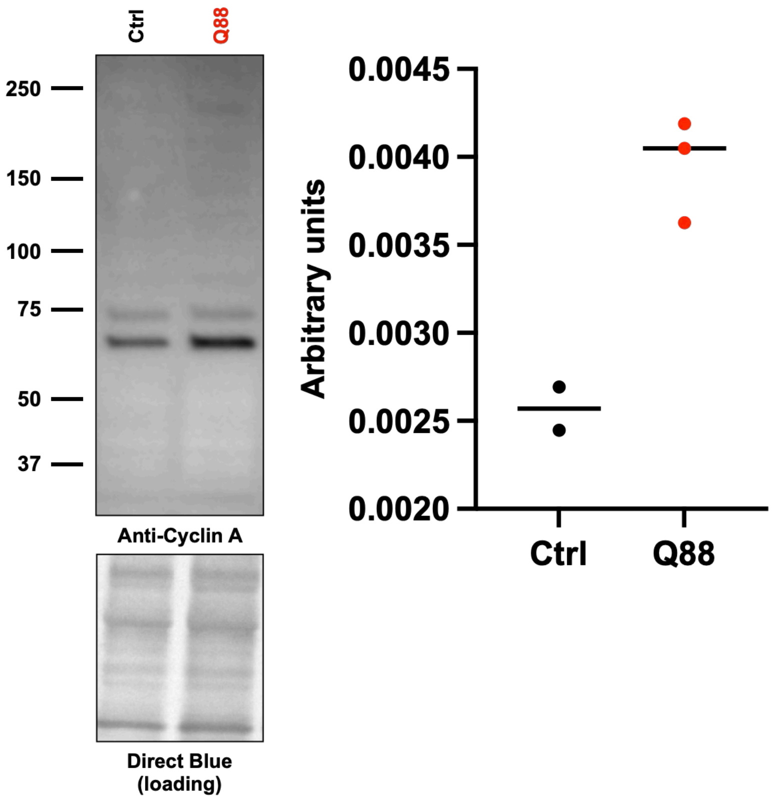

**Supplemental figure 10**

```
########################################
# Program: needle
# Rundate: Sun 11 Aug 2024 12:29:31
# Commandline: needle
#    -auto
#    -stdout
#    -asequence emboss_needle-I20240811-122926-0499-37714441-p1m.asequence
#    -bsequence emboss_needle-I20240811-122926-0499-37714441-p1m.bsequence
#    -datafile EBLOSUM62
#    -gapopen 10.0
#    -gapextend 0.5
#    -endopen 10.0
#    -endextend 0.5
#    -aformat3 pair
#    -sprotein1
#    -sprotein2
# Align_format: pair
# Report_file: stdout
########################################

#=======================================
#
# Aligned_sequences: 2
# 1: Grunge
# 2: ATN1(Q7)
# Matrix: EBLOSUM62
# Gap_penalty: 10.0
# Extend_penalty: 0.5
#
# Length: 2216
# Identity:     385/2216 (17.4%)
# Similarity:   539/2216 (24.3%)
# Gaps:        1247/2216 (56.3%)
# Score: 870.0
#
#
#=======================================

Grunge             1 MAASTQGEIRVGPGHQVNDVYAKLPDYNPISSFPIDKETDERELEESRWS     50

ATN1(Q7)           1 --------------------------------------------------      0

Grunge            51 PGVVADGDLLMFLRAARSMAAFQGMCDGGLEDGCLAASRDDTTINALDVL    100

ATN1(Q7)           1 --------------------------------------------------      0

Grunge           101 HDSGYDPGKALQALVKCPVSKGIDKKWTEDETKKFIKGLRQFGKNFFRIH    150

ATN1(Q7)           1 --------------------------------------------------      0

Grunge           151 KDLLPHKDTPELVEFYYLWKKTPGANNNRPHRRRRQSALRRNRVTRANNS    200

ATN1(Q7)           1 --------------------------------------------------      0

Grunge           201 NSNTPPKKEDTPEPQTATTATAAATAASETASRSSPAVSKEENSSLTEDD    250

ATN1(Q7)           1 --------------------------------------------------      0

Grunge           251 ASECDSDSSLTHKRDESPSRMRTRNKQQNNNSSTSSGNNTAGNGGGNATS    300

ATN1(Q7)           1 --------------------------------------------------      0

Grunge           301 ISSGSTGGGAAGGNSSSKDQSANAVANGKRPKRGSETPDVSGGASVDSPK    350

ATN1(Q7)           1 --------------------------------------------------      0

Grunge           351 TPTKAVAESSANKRKGGKQDTPNKKKRTEQESNEPSAHEENAIKEKRKRP    400

ATN1(Q7)           1 --------------------------------------------------      0

Grunge           401 DSPVESMNSDSRPDSVLDDGESNTTDTTTAEQQSTKDSKETVSCKEEREM    450
                                                     ..|:.:|:::|.:..|:.
ATN1(Q7)           1 --------------------------------MKTRQNKDSMSMRSGRKK     18

Grunge           451 VTNDLEAKAEEKAIKAEALAEDSKDSAIKNMDEETNIQAPSSADTSLVDG    500
                          ||....:.:::...|                  :|....||..||
ATN1(Q7)          19 -----EAPGPREELRSRGRA------------------SPGGVSTSSSDG     45

Grunge           501 PNPNALPSPVAAPITMKVPTIATVEALNASVDRKEAIEKMESCDSDPEML    550
                                                .|...|:.|.:......|.|::.
ATN1(Q7)          46 ---------------------------KAEKSRQTAKKARVEEASTPKVN     68

Grunge           551 KKLATIKQEVSPQQQQHMQQQSQQQMQQQLAPVGIPQPPSCPPSESVYIK    600
                     |:..:  :|:|..:.:......:.:.:|:|     |:|.|  ||:.
ATN1(Q7)          69 KQGRS--EEISESESEETNAPKKTKTEQEL-----PRPQS--PSDL----    105

Grunge           601 KEPMEDSMDATC--NQNSNEPQDL-----KVKIEIKNEDALKH---SAGG    640
                          ||:|...  :..|::|:|:     .....|.:..::::   |:.|
ATN1(Q7)         106 -----DSLDGRSLNDDGSSDPRDIDQDNRSTSPSIYSPGSVENDSDSSSG    150

Grunge           641 LPPSGPCAP--PSALHPLSGAPVESGQEPLHLQHMPHGQVTTQPPPGYLI    688
                     | ..||..|  |..|.|.|..|.:|..........||..||   |.||
ATN1(Q7)         151 L-SQGPARPYHPPPLFPPSPQPPDSTPRQPEASFEPHPSVT---PTGY--    194

Grunge           689 DGQLKYGP----------SGQGVPPQPPQLHSDAAGGVSGAPP-------    721
                          :.|          :..|.||..|||:....|||...||
ATN1(Q7)         195 -----HAPMEPPTSRMFQAPPGAPPPHPQLYPGGTGGVLSGPPMGPKGGG    239

Grunge           722 -----GAPTTPQKYPPEMEMKFAPQDLKYPPPPPLDALKYSQEMQAAAAA    766
                          |.|...:::|              ||..|:           :.::
ATN1(Q7)         240 AASSVGGPNGGKQHP--------------PPTTPI-----------SVSS    264

Grunge           767 AAAAGKYDMKYMMEQQGKYNVELSAAHQPPSKPGYQDSLKIPDIKPGFGH    816
                     :.|:|                      .||:||        |....|.|:
ATN1(Q7)         265 SGASG----------------------APPTKP--------PTTPVGGGN    284

Grunge           817 LPHNVGSPLDAAHKYGPPPTSQESQQQQPQPPAHQVPPGATPPPGIAMPK    866
                     ||.            .|||.:           ...|.|...|||.:
ATN1(Q7)         285 LPS------------APPPAN-----------FPHVTPNLPPPPAL----    307

Grunge           867 PHYQHDVQTPPLGRPFEPTGLMLKYGDPLAAKYGPPQDLKYPMPPVSQAG    916
                                  ||.
ATN1(Q7)         308 -------------RPL----------------------------------    310

Grunge           917 PADVKPYGGENLIKSSPYGPPPESPIDASARSTPGQDSQGSNSNSQPPSM    966
                                                             :|:::.||.:
ATN1(Q7)         311 ----------------------------------------NNASASPPGL    320

Grunge           967 PPQPQQFQSPHPSPHMPSPAGGGLPPGMHPQNLIHGPPPGAAGGSGPQPP   1016
                     ..||  .....||||......||||||  |:             .||...
ATN1(Q7)         321 GAQP--LPGHLPSPHAMGQGMGGLPPG--PE-------------KGPTLA    353

Grunge          1017 PPPTSLHQPTPTSAGPPSLQHGLHPGHQHSQLSVASSIPPSSIGIPPTLS   1066
                     |.|.|| .|..:||..|.::.........|..:.:||...||...|...|
ATN1(Q7)         354 PSPHSL-PPASSSAPAPPMRFPYSSSSSSSAAASSSSSSSSSSASPFPAS    402

Grunge          1067 TMAPSHMHPHLHPHAHLQGLHRPHDLPPSMHPHAPMPLSLQGHPQHGHGL   1116
                     ...||:                ||..||      |..||:...
ATN1(Q7)         403 QALPSY----------------PHSFPP------PTSLSVSNQ-------    423

Grunge          1117 PPSHTSQQQQQQQQQQQPGGPAGTVRTPSPAQQPPRSMHDPQSSREPPTS   1166
                     ||.:|           ||..|:..|.:..|...||.......|:..|...
ATN1(Q7)         424 PPKYT-----------QPSLPSQAVWSQGPPPPPPYGRLLANSNAHPGPF    462

Grunge          1167 QPSTTMAGSSGPGGPPPQQSPHAHRTSPLPGLAG-SGPPPPGLIGHPMAI   1215
                     .|||   |:.....||.....|.|:........| |||||||...||
ATN1(Q7)         463 PPST---GAQSTAHPPVSTHHHHHQQQQQQQHHGNSGPPPPGAFPHP---    506

Grunge          1216 HPHLAHLPPGHPAHAALAHPGHHLLSHSIAGLGPGGGPIALLAGPGGLGG   1265
                           |..|...|   |||  :.:|.|:..|.|        ..||
ATN1(Q7)         507 ------LEGGSSHH---AHP--YAMSPSLGSLRP--------YPPG----    533

Grunge          1266 IPESALSRRTPPSHLPHSHASSAPLTAHSVASMTSTSMSLTTSTVPSSAF   1315
                                |:|||..|:
ATN1(Q7)         534 -----------PAHLPPPHS------------------------------    542

Grunge          1316 SRASPSVQISSSGGGPSGPGSVGPGGMPNSSAAAAAAAAAHRAASPASSV   1365
                            |:|.|..||:||                          |.||.
ATN1(Q7)         543 -------QVSYSQAGPNGP--------------------------PVSSS    559

Grunge          1366 SSLSRQSPLHPVPQSPLSHHPSSSALSAAAAAVAERDRHALMRQQSPHMT   1415
                     |:.|..:.....|.|    |||.|                    |.|...
ATN1(Q7)         560 SNSSSSTSQGSYPCS----HPSPS--------------------QGPQGA    585

Grunge          1416 P---PPVSNASLMASPLSKMYAPQPGQRGLGTSPPPHLRPGASPPVIRHP   1462
                     |   |||...:..::.||.:.|
ATN1(Q7)         586 PYPFPPVPTVTTSSATLSTVIA----------------------------    607

Grunge          1463 QMPLPLPLIAPGGGIPQIGVHPGQSPYPHPLLHPSVFYSPHHHPFNSPYG   1512
                                                       :|..||..:...||
ATN1(Q7)         608 ----------------------------------TVASSPAGYKTASP--    621

Grunge          1513 YAPYGPGFPAYMKPPPQPGQLDPAAVMAAHHAGLQGPPPQQMRQDEQNAA   1562
                          ||.|.|.|..|.||....|.
ATN1(Q7)         622 -----PGPPPYGKRAPSPGAYKTAT-------------------------    641

Grunge          1563 AAAAQAAAEKQHQAAAAAAAQQHKAPQQQQPGGMPPNKPPTPKTPQGPG-   1611
                                                   |.|..|..||:.:|...||
ATN1(Q7)         642 ------------------------------PPGYKPGSPPSFRTGTPPGY    661

Grunge          1612 -GGMPPGMGG---PGTPTGLPPGAYPGSHMPGYPQGPPHGSPFAPQDGQP   1657
                      |..||...|   ||:|| :.||..|.:...|.|..||  .|.||..|.|
ATN1(Q7)         662 RGTSPPAGPGTFKPGSPT-VGPGPLPPAGPSGLPSLPP--PPAAPASGPP    708

Grunge          1658 HGLKPTSHMDALRAHAHSANSAGMGGGHHPTEPLPIDIEPDPEPEIPSPT   1707
                       |..|                              .|:.:|..|..:|.
ATN1(Q7)         709 --LSAT------------------------------QIKQEPAEEYETPE    726

Grunge          1708 HNIP--RGPSPEAKPDDTECHRSQSAIFVRHIDRGDYNSCTRTDLIFKPV   1755
                     ..:|  |.|||..|..|...|.||||.|.:|:||| :|||.|:||.|.|:
ATN1(Q7)         727 SPVPPARSPSPPPKVVDVPSHASQSARFNKHLDRG-FNSCARSDLYFVPL    775

Grunge          1756 ADSKLARK---------REERDRKLAEKERERRQQQQQQQQQQQQQQAAA   1796
                     ..||||:|         ||...|...||||||.::::::::::::::...
ATN1(Q7)         776 EGSKLAKKRADLVEKVRREAEQRAREEKEREREREREKEREREKERELER    825

Grunge          1797 AQQAAQQAKMKAELK--------------------PPY--ADTPALRQLS   1824
                     :.:.||:.:...|..                    |||  .||||||.||
ATN1(Q7)         826 SVKLAQEGRAPVECPSLGPVPHRPPFEPGSAVATVPPYLGPDTPALRTLS    875

Grunge          1825 EYARPHVAFSPVEQMVPYHHPMG------------------PMYRERELE   1856
                     ||||||| .||..:..|::.|:|                  |..||||.|
ATN1(Q7)         876 EYARPHV-MSPGNRNHPFYVPLGAVDPGLLGYNVPALYSSDPAARERERE    924

Grunge          1857 --------------EIKNAQAAAASQSRLDP-H-----WMEYYRR-----   1881
                                   |:|        .|.|:| |     .::.:.|
ATN1(Q7)         925 ARERDLRDRLKPGFEVK--------PSELEPLHGVPGPGLDPFPRHGGLA    966

Grunge          1882 ------GIHPSQFPLYANPAISQMERERLGI-------PPPHH-------   1911
                           |:||  ||.:  |::..:|||||.:       |...:
ATN1(Q7)         967 LQPGPPGLHP--FPFH--PSLGPLERERLALAAGPALRPDMSYAERLAAE   1012

Grunge          1912 ---------VGLDP--GEHMIRLTREYHAHSHTHLHLPLHPQPQPPEAGF   1950
                              :|.||  ...|:.:|..:|.|||.|.||.||.|.....|..
ATN1(Q7)        1013 RQHAERVAALGNDPLARLQMLNVTPHHHQHSHIHSHLHLHQQDAIHAASA   1062

Grunge          1951 QLPPNVG-----------QYPR---PNMLIPREPHSDVLLR---------   1977
                     .:.|.:.           .||.   ||.|:|...|.:.:||
ATN1(Q7)        1063 SVHPLIDPLASGSHLTRIPYPAGTLPNPLLPHPLHENEVLRHQLFAAPYR   1112

Grunge          1978 ---------MSYADQLQ--YLQAAEFQRQSLHDQYFRNQLR---------   2007
                              ||.|.|||  :.|:||.||.:|..|.:.:...
ATN1(Q7)        1113 DLPASLSAPMSAAHQLQAMHAQSAELQRLALEQQQWLHAHHPLHSVPLPA   1162

Grunge          2008 ----------------   2007

ATN1(Q7)        1163 QEDYYSHLKKESDKPL   1178


#---------------------------------------
#---------------------------------------
```
